# Computational design of sequence-specific DNA-binding proteins

**DOI:** 10.1101/2023.09.20.558720

**Authors:** Cameron J. Glasscock, Robert Pecoraro, Ryan McHugh, Lindsey A. Doyle, Wei Chen, Olivier Boivin, Beau Lonnquist, Emily Na, Yuliya Politanska, Hugh K. Haddox, David Cox, Christoffer Norn, Brian Coventry, Inna Goreshnik, Dionne Vafeados, Gyu Rie Lee, Raluca Gordan, Barry L. Stoddard, Frank DiMaio, David Baker

**Affiliations:** Department of Biochemistry, University of Washington, Seattle, WA, USA; Institute for Protein Design, University of Washington, Seattle, WA, USA; Department of Physics, University of Washington, Seattle, WA, USA; Division of Basic Sciences, Fred Hutchinson Cancer Center, Seattle, Washington, USA; Program in Genetics and Genomic, Duke University, Durham, NC, USA; Center for Advanced Genomic Technologies, Duke University, Durham, NC, USA; Department of Bioengineering, University of Washington, Seattle, WA, USA; Department of Biochemistry, Stanford University School of Medicine, Palo Alto, CA USA; Department of Medicine, Division of Hematology, Stanford University, Stanford, CA, USA; BioInnovation Institute, DK2200 Copenhagen N, Denmark; Howard Hughes Medical Institute, University of Washington, Seattle, WA USA; Department of Biostatistics and Bioinformatics, Department of Computer Science, Department of Molecular Genetics and Microbiology, Duke University, Durham, NC, USA

**Author notes:** These authors contributed equally to this work (Cameron J. Glasscock, Robert Pecoraro, Ryan McHugh).

## Abstract

Sequence-specific DNA-binding proteins (DBPs) play critical roles in biology and biotechnology, and there has been considerable interest in the engineering of DBPs with new or altered specificities for genome editing and other applications. While there has been some success in reprogramming naturally occurring DBPs using selection methods, the computational design of new DBPs that recognize arbitrary target sites remains an outstanding challenge. We describe a computational method for the design of small DBPs that recognize specific target sequences through interactions with bases in the major groove, and employ this method in conjunction with experimental screening to generate binders for 5 distinct DNA targets. These binders exhibit specificity closely matching the computational models for the target DNA sequences at as many as 6 base positions and affinities as low as 30–100 nM. The crystal structure of a designed DBP-target site complex is in close agreement with the design model, highlighting the accuracy of the design method. The designed DBPs function in both *Escherichia coli* and mammalian cells to repress and activate transcription of neighboring genes. Our method is a substantial step towards a general route to small and hence readily deliverable sequence-specific DBPs for gene regulation and editing.

## Main text

Nature employs a wide diversity of DNA binding protein (DBP) domains for targeting specific sequences(*1*) which are often structurally coupled to each other and to effector regions conferring enzymatic, binding and regulatory functions(*2*, *3*). Despite intensive study, the DNA binding affinity and specificity of natural proteins remain difficult to predict(*4*), and the high free energetic cost of desolvating the highly polar DNA surface presents a challenge to the *de novo* design of new DBPs. For these reasons, while computational *de novo* design has had considerable recent success in generating binders to arbitrary protein structures(*5*), mostly at hydrophobic patches, computational approaches for DBP engineering have thus far been limited to redesigning interfaces of existing native protein-DNA complex structures(*6–10*). These efforts have been constrained by the rigid geometry of the starting scaffold shape and orientation relative to DNA(*11*), which restrict the possible target sequences that can be recognized(*12*). A general solution to generating compact, customizable DBPs would have significant advantages in modularity and deliverability and be highly complementary to the state-of-the-art in gene regulation, gene editing, and nucleic acid diagnostics which primarily employ Cys_2_His_2_ zinc finger (ZF) domains(*13*, *14*), transcription activator-like effectors (TALEs)(*15*, *16*), and CRISPR-Cas(*17*). While these tools have proven powerful, each has limitations: ZFs can be laborious to engineer, and the size of TALE and CRISPR-Cas systems complicates their delivery in therapeutic applications; CRISPR-Cas systems also require an extra guide RNA component and target sites are constrained by protospacer adjacent motif (PAM) requirements(*17*). These systems will undoubtedly continue to be improved, but their fixed overall backbone topologies can constrain interaction specificity and close integration with diverse effector domains.

### Design strategy

We reasoned that it would be possible to achieve general DNA sequence recognition using small compact proteins by sampling a wide variety of structures and binding modes to find those which are optimal for targeting specific sequences of interest. We previously developed a general method for designing specific protein binders to arbitrary protein targets based on this concept(*5*), but sequence-specific DNA binding requires overcoming several additional challenges. First, binding the DNA double helix, with major and minor grooves, requires sufficient shape complementarity with the DNA backbone to precisely position specific amino acid residues to interact with the DNA base edges. Second, recognition of DNA sequences requires distinguishing between the subtle changes in individual atom placements among the four bases(*18–24*) which alter the landscape of potential molecular contacts. Third, in contrast to designed protein-protein contacts mostly mediated by orientation-agnostic hydrophobic patches(*5*), the majority of accessible DNA base atoms require hydrogen bond interactions with polar sidechains for specific recognition(*25*). Not only are polar interactions harder to model accurately, but the longer polar sidechains have considerable conformational flexibility, making structure modeling more difficult and increasing opportunities for off-target base interactions through alternate sidechain rotamer conformations. To address these challenges, we formulated a set of design principles (**Fig. 1A**) and sought to develop a design pipeline implementing them (**Fig. 1, B to G**).

**Fig. 1.**
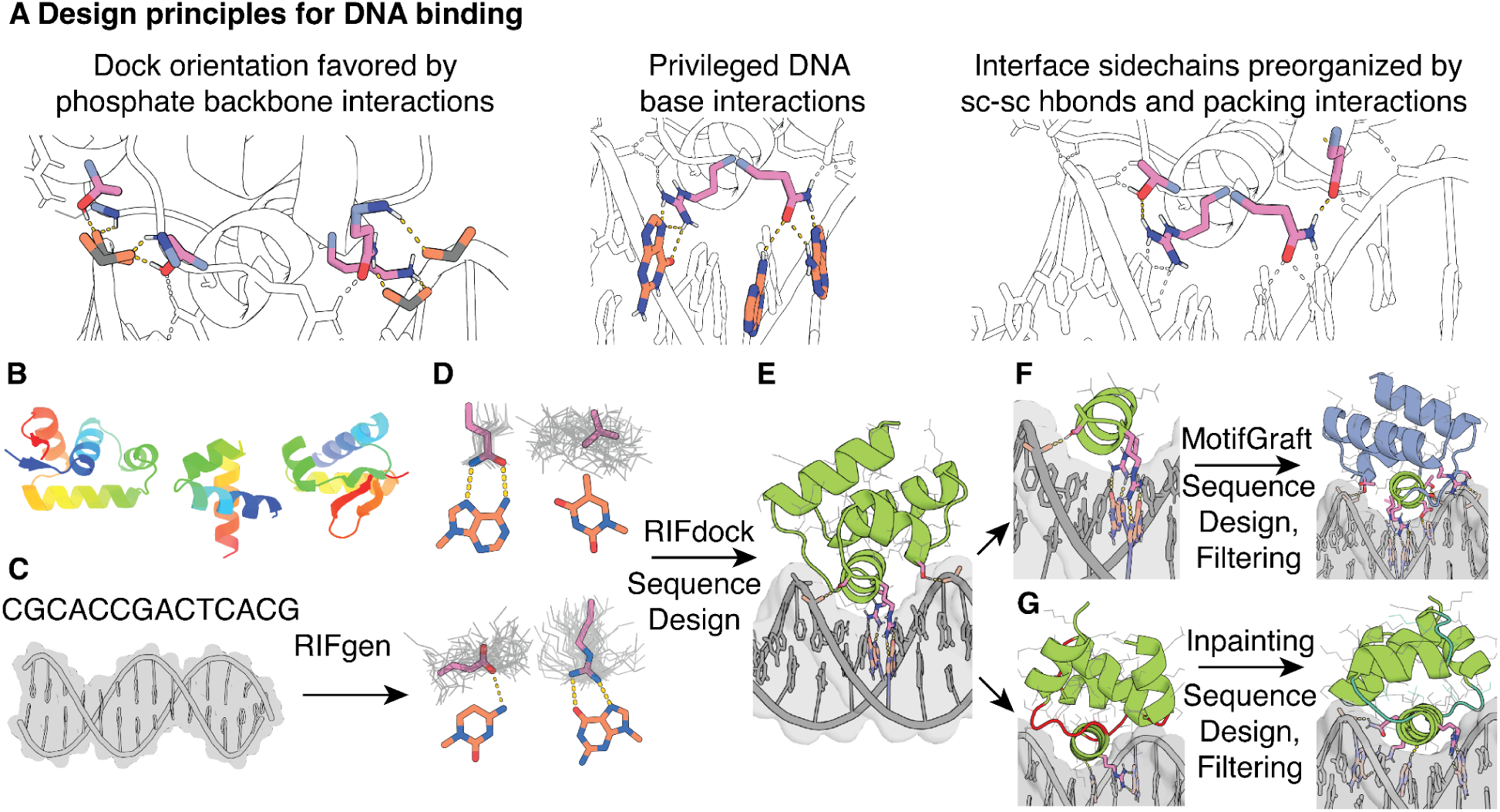
**Overview of the DNA binder design pipeline.** (**A**) Design principles for design of sequence-specific DBPs. (**B**) HTH backbone scaffold library generated from metagenomic sequences. (**C**) DNA target, starting with either a specific nucleotide sequence modeled as B-DNA or a DNA crystal structure. (**D**) Generation of RIF (gray) to form base-specific hydrogen bonds and hydrophobic packing interactions. Example rotamers (pink) are generated for nucleotide bases (orange; clockwise from upper left: adenine, thymine, guanine, cytosine). (**E**) Docking of scaffolds onto the RIF to identify seed interactions and placements with base-specific contacts, followed by sequence optimization of the DNA-scaffold interactions using Rosetta or LigandMPNN-based sequence design and Rosetta modeling. (**F**) Recognition helices making multiple favorable interactions with the target are extracted from first round designs, and grafted onto the scaffold library, followed by further rounds of interface sequence design and filtering for favorable interactions. (**G**) Inpainting of the protein loops (red) results in new connecting loops (teal) between the helical portions of the design, followed by further rounds of interface sequence design and filtering.

For design challenge one, we hypothesized that interactions with the phosphate backbone, such as the backbone amide-mediated hydrogen bond interactions with DNA phosphate oxygens (hereon called mainchain-phosphate hydrogen bonds) that are frequently observed in native DBP structures, could enable precise placement of designed scaffolds such that residues designed to make specific base contacts interact with these in the intended geometry. Hence we reasoned that DNA binding would require a custom scaffold library to satisfy the hydrogen bond requirements of the DNA backbone phosphates and the DNA bases, which substantially constrains viable scaffold geometries. To generate such a library of small (< 65 amino acid) and structurally diverse scaffolds, we took advantage of the vast amount of metagenome sequence data and the accuracy of deep learning based protein structure prediction (**fig. S1**). We carried out sequence searches for helix-turn-helix (HTH) DNA-binding domains(*26*), generated AlphaFold2 (AF2) structure predictions(*27*), and filtered these based on prediction confidence (pLDDT) and TMscore to known HTH domain structures(*28*). This resulted in a library of ∼26,000 HTH scaffolds which finely sample different helix orientations and loop geometries (**fig. S1**) (see Methods).

We docked the scaffolds against specific DNA target structures seeking to maximize the potential for specific sidechain-base interactions (**Fig. 1, B to D)**. To do this, we extended the RIFdock approach(*5*) to protein-DNA interactions (see Methods), which finely samples many possible *de novo* docks for each scaffold. RIFdock begins by enumerating a large and comprehensive set of disembodied sidechain interactions, called a Rotamer Interaction Field (RIF), that make favorable interactions with the desired target. We focused RIF generation on polar and nonpolar interactions with nucleotide base atoms in the major groove of the DNA target, with an emphasis on protein sidechain-DNA base hydrogen bonding interactions that are statistically more probable in native protein-DNA complexes(*29*). In RIFdock, we constrained the RIF DNA base-specific interactions to the HTH recognition helix, and obtained ten million distinct docks for sequence design. Docking with HTH scaffolds allowed us to find placements with both mainchain-phosphate hydrogen bonds and base-contacting RIF sidechains, resolving the first design challenge.

To address design challenge two – recognizing specific DNA bases – we used either Rosetta-based sequence design or an extended version of the deep learning–based ProteinMPNN sequence design software to promote folding of the scaffold and high affinity binding to the DNA target (**Fig. 1E**, see Methods). As originally described, the ProteinMPNN graphical model generates amino acid sequences purely based on protein backbone coordinates, but a recent extension to incorporate ligand and DNA atoms in the interaction graph, called LigandMPNN, enables design in the presence of specific DNA target sites. While the Rosetta-based sequence design protocol was constrained by a position-specific scoring matrix (PSSM) for each scaffold, LigandMPNN was not constrained by such information so that sequence design across the interface and within the binder was purely based on the structure of the designed complex. To reduce the computational cost of full sequence design on the millions of generated scaffold docks for each target site, we first repacked only the RIF sidechain residues in the context of the target to remove potential clashes between designed sidechains. Docks for which good protein-DNA interactions could be achieved without sidechain clashes were then subjected to multiple iterations of full sequence design, alternating with Rosetta backbone relaxation to maximize complementarity to the target sequence. We generated 200,000–300,000 designed complexes per target. From this large set of designs, we selected those with the most favorable free energy of binding (Rosetta ΔΔG), contact molecular surface area(*5*) and interface hydrogen bonds, the fewest interface buried unsatisfied hydrogen bond donors and acceptors, and with bidentate sidechain-base hydrogen bonding arrangements frequent in the Protein Data Bank (PDB) (see Methods for full details).

To address design challenge three – precise geometric sidechain placement – we hypothesized that specificity and affinity would be improved in designs with highly preorganized interface sidechains. We reasoned that preorganization would be especially important for long polar sidechains with many possible conformations. We achieved preorganization through sidechain (sc)-sc hydrogen bonding and assessed it using the Rosetta RotamerBoltzmann calculation(*30*). By selecting only designs with native-like preorganization of key contacts (**fig. S2**), we aimed to achieve the level of precision required for specific DNA binding.

Following selection based on the above criteria, and clustering by sequence identity, the monomeric structures of the hundreds to thousands of designs which remained for each target were predicted based on their sequences using AF2, and designs that deviated from their original design models were discarded. The remaining predicted monomer structures were superimposed onto the design complex by alignment on the interface residues of the original design and relaxed with Rosetta in the context of the DNA. Designs with the most favorable DNA binding interactions post-superimposition, as assessed with the above metrics, were selected for experimental characterization. To obtain additional high-quality designs, the DNA interacting segments of the filtered designs were extracted, clustered, and grafted back into the original *in silico* scaffold library, followed by a second round of sequence design (**Fig. 1F)**(*5*). We also diversified the best designs using RoseTTAFold Inpainting(*31*) focused on the resampling of scaffold loops followed by sequence design (**Fig. 1G)**. We generated at least 10,000 designs for each DNA target that passed all the structural and DNA interaction filters using a combination of these approaches.

### Design generation and screening with yeast display cell sorting and deep sequencing

We created three sets of designs using variations of the overall design approach. In the first set, we generated 21,488 designs using Rosetta-based sequence design, the motif grafting strategy, and our custom scaffold library of AF2-predicted native DNA-binding domains. In this set, the double-stranded DNA (dsDNA) targets were the DNA portions of co-crystal structures. In the second design set, we generated 12,273 designs against the same DNA sequences, with the LigandMPNN sequence design strategy and the motif grafting approach for backbone resampling. In this case, rather than designing only against the dsDNA conformations found in each target’s respective crystal structure, we also designed against straight B-DNA of the same sequences (6,608 designs B-form, 5,666 crystal-derived). The LigandMPNN approach was less effective at generating designs with high contact molecular surface, likely because of the ability of Rosetta to relax the protein backbone during sequence design, but ultimately produced designs with more favorable free energy of binding (Rosetta ΔΔG) and an increased number of hydrogen bonds to bases (**fig. S3**). Finally, in the third set we generated 100,000 designs using the LigandMPNN-based design pipeline and the inpainting-based backbone remodeling protocol against 11 unique B-DNA targets.

For each set of designs, synthetic oligonucleotides (230 base pairs) encoding the 50–65-residue designed proteins were ordered in a single pool and cloned into a yeast surface-expression vector. Cells containing designs that bound each DNA target were enriched by several rounds of fluorescence-activated cell sorting (FACS) using fluorescently labeled target dsDNA oligos. The naive and sorted populations for each DNA target were deep sequenced, and the frequency of each design in the starting population and after each sort was determined. From this analysis, we identified 97 designs that were substantially enriched (>100x) in pools sorted with their intended dsDNA target compared to the naive library.

We tested these 97 designs as individual clones in a 96-well screening format and found detectable binding for 44 of them (**fig. S4**). The remainder may result from doublet transformants in the yeast pool or are very weak binders that were enriched under higher dsDNA oligo concentrations. For each of the 44 successful designs, we knocked out the DNA binding interface by substituting the 2–3 residues making the most extensive interactions with the DNA bases such that the AF2 Cα RMSD was < 2 Å to the original design model (**table S1**). These knockout mutations completely or substantially disrupted binding for all designs that had detectable binding on yeast (**fig. S4**), indicating that the functional designs are working as intended.

### Design conformation and DNA-side footprinting of binding specificity

We performed an all-by-all screen of DBP design hits to 13 unique dsDNA targets **(fig. S5, table S2**). Several designs exhibited a strong preference for only their designed target sequence (e.g. DBPs 6, 9, 62), others exhibited a strong preference for 2 or 3 of the sequence targets (e.g. DBPs 1, 52, 60), and a few bound to most of the targets (e.g. DBPs 23, 44, 89). To try to understand these observed binding preferences, each tested DNA sequence was threaded onto each design complex model at all possible basepair alignments, the alternative complex models were relaxed with Rosetta, and the model with the most favorable Rosetta ΔΔG was selected. We found a modest correlation between the predicted free energy of binding and the extent of off-target binding (**fig. S5**); for DBPs 44 and 89, Rosetta ΔΔGs comparable to the original targeted sequence were obtained for most of the off target sites, consistent with the observed low specificity. Overall, we found that 14 designs bound with specificity closely consistent with the design models (DBPs 5, 6, 9, 35, 43, 69, 47, 48, 51, 56, 57, 60, 62, 85), including binders for 5 unique DNA sequences (Sequences A–E).

We used a yeast display competition assay to characterize the DNA binding site specificity of a subset of the designs (**Fig. 2, A to E, left and fig. S6**). Addition of non-biotinylated competitor dsDNA to biotinylated target sequence reduced binding signal by flow cytometry, and scanning base substitutions through the competitor revealed positions important for binding (**Fig. 2, A to E, middle**). DBPs 6, 35, 48, 56, and 62 exhibited specificities consistent with the designed sidechain-base interactions. For example, in DBP6, R31 and R36 in the design model form bidentate hydrogen bonds with the guanines of base pair positions G12 and C9, respectively, while T32 forms a hydrogen bond with C10. Substitution of the bases at positions 9, 10, and 12 eliminated competition, indicating specificity for the GCxG motif as expected (**Fig. 2A**). DBP62 exhibited specificity for its target site despite having relatively few base-specific hydrogen bonding interactions; specificity in this case may result from the very tightly packed interface (**Fig. 2E**).

**Fig. 2.**
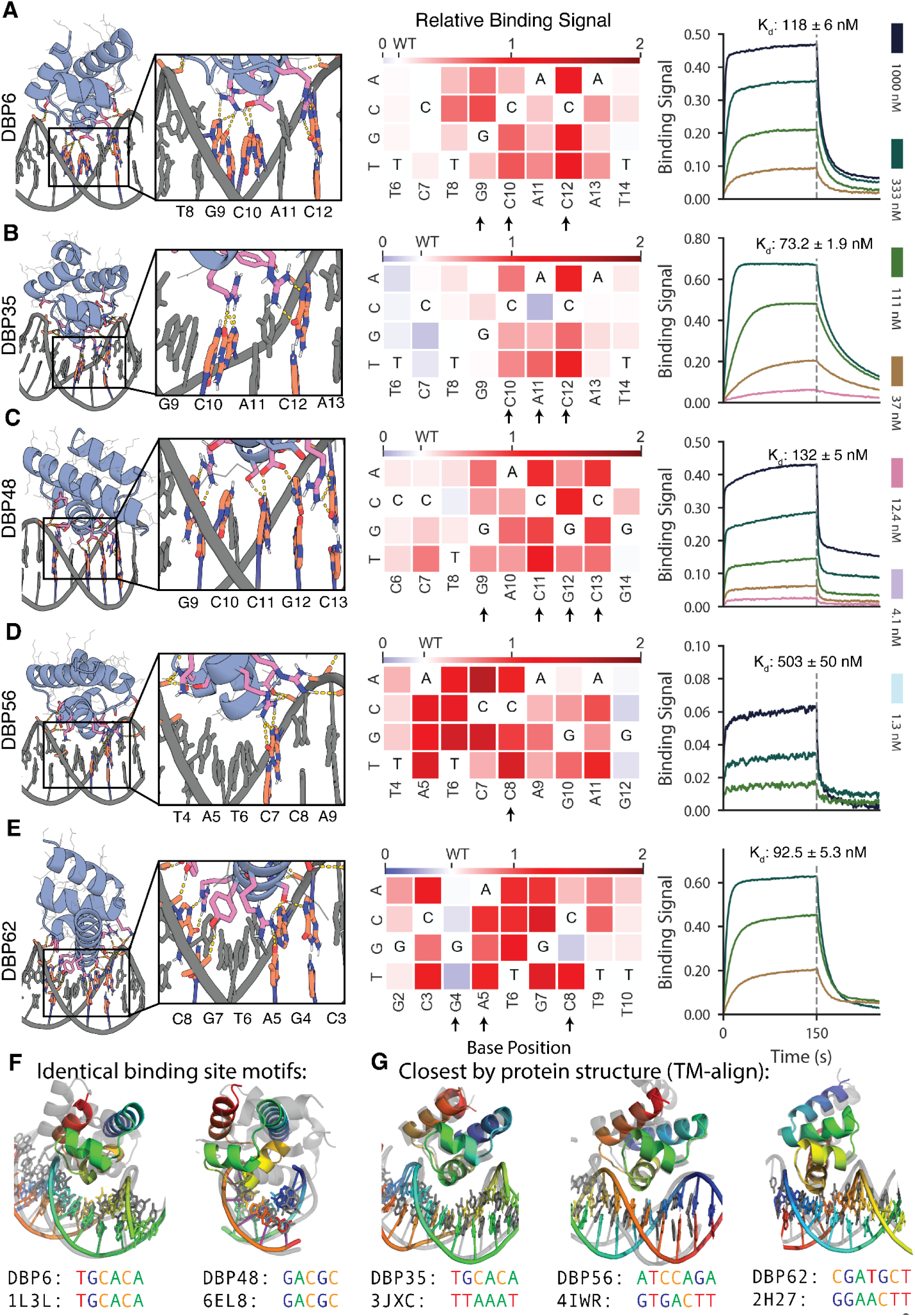
Designed DBPs bind with high affinity and specificity to their intended target sites. **(A to E)** Characterization of DBPs 6, 35, 48, 56, and 62, respectively. **Left,** Computational design models of characterized designs at the DNA-binding interface. DNA bases and protein residues involved in hydrogen bonding interactions are shown in orange and pink, respectively. Hydrogen bonds are highlighted with dashed yellow lines. **Middle,** Relative binding activity (PE/FITC normalized to the no-competitor condition) from flow cytometry analysis in yeast display competition assays with all possible DNA base mutations at each position of the competitor oligo. Blue indicates competitor mutations where competition was stronger than with the wild-type competitor, while red indicates competitor mutations where competition was weaker. Arrows indicate base positions contacted with hydrogen bonds or hydrophobic contacts to base atoms in the design model. Additional characterized designs are shown in **fig. S6**. **Right,** Binding of purified miniprotein designs to the DNA target with BLI. Each line represents biotinylated dsDNA target dilutions by ⅓. The highest DNA target concentration is indicated in each plot. Additional characterized designs are shown in **fig. S8**. (**F**) DBPs 6 and 48 (colored) differ in both structure and docking mode to native co-complex structures with matching DNA binding sites (gray). (**G**) DBP35 has similar structure and dock to the closest match in the PDB, but binds a distinct DNA target site, while DBPs 56 and 62 have structures similar to the closest matches but different docks and DNA target sites. DBP48 was analyzed with sequence C due to its improved binding signal and nearly identical modeled binding sites (**fig. S5)**; all other designs were analyzed with their designed target sequence.

Genes encoding the designs were encoded for *E. coli* expression and purified proteins were evaluated for binding *in vitro*. Most of the selected designs were in the soluble fraction, readily purified by Ni^2+^-NTA chromatography, and appeared monodisperse by size exclusion chromatography (**fig. S7**). Binding to the biotinylated dsDNA oligo was assessed using biolayer interferometry, and all designs were found to bind with binding affinities ranging from 30–500 nM (**Fig. 2, A to E, right and fig. S8**).

Although some of the designs were targeted towards DNA sequences found in crystal structures, the designed DBPs and their sequence preferences are novel as assessed by comparison of the binding site motifs to co-complex structures of native DBPs in the PDB containing a protein helix in contact with bases in the DNA major groove. We found that some designs (DBPs 6, 35, 48) preferred a similar motif as native DBP structures but had substantially unique interfaces and docking orientations, while other designs (DBPs 56, 62) bound novel sequences found neither in the PDB (**Fig. 2F and fig. S9, A to C**) nor in the JASPAR non-redundant transcription binding profile database(*32*, *33*) (**fig. S10**).

Our binder design method aims to effectively sample diverse scaffold-DNA docks to find solutions optimal for binding the target DNA sequence. The method could, in principle, recover solutions similar to known native DBP-DNA complexes. To investigate this, we compared the structures of our designed DBPs to native DBP domains in DNA co-crystal structures in the PDB by TM-align(*28*) (closest structures: **Fig. 2G; fig. S9, D to H; and table S3**). We found that the overall folds of the designed scaffolds had matches in the PDB, but the placement of the scaffold relative to the DNA generally differed, as expected given the *de novo* docking step in our approach. None of the closest matches by protein structure had more than 3 out of 7 common bases at the aligned DNA binding site positions and the sidechain-base hydrogen bond networks differed substantially. For all designs that bound their DNA targets, we also performed Blastp searches of the non-redundant protein sequences database (nr)(*34*) and found that most had sequence similarity to native metagenome protein sequences ranging from 40–60% (**table S1).** Overall, these analyses suggest that our approach was able to utilize and expand upon the known native docking space, while exploring new sequence space, to identify effective DBP designs against the specified target sequences.

To evaluate the importance of backbone sampling through docking, we examined the ability of LigandMPNN-based sequence design to generate interfaces passing our *in silico* metrics when starting from crystal structures of native co-complexes rather than *de novo* docks. Starting from co-crystal structures with high TM-align scores to the designed DBPs, we mutated the DNA sequence *in silico* to the target sequence, and redesigned the sequence using LigandMPNN. We found that designs based on fixed native backbones failed to recover most of the base-specific hydrogen bonds present in the designs produced by our docking pipeline (**fig. S9I**). In the few cases where native redesign did recover multiple base-specific hydrogen bonds, such as DBPs 6 and 35, the *de novo*-docked design models scored better on sidechain preorganization by the RotamerBoltzmann metric (**fig. S9J**), suggesting non-hydrogen bond features of the interface that may be critical for specific binding and require precise docking configurations. Overall, our design method is able to sample and identify designs that would not be identified through structure-based redesign of native protein-DNA co-complexes and generate specific binders for unique DNA sequences that are not known to be recognized by native proteins.

### Structural validation by x-ray co-crystallography and footprinting of the protein binding surface

We solved the co-crystal structure of DBP48 in complex with its preferred target sequence, and found very close agreement to the design model (**Fig. 3A and fig. S11)**. Cα-RMSDs of the co-crystal structure to the design model were 0.641 Å for the binder alone, and 1.907 Å all-atom RMSD across the protein-DNA complex. Among interface residues forming key interactions with bases, R38 and S39 were in the closest agreement and formed the expected sidechain-base hydrogen bonds (**Fig. 3B)**. D43 and R49 did not form the expected hydrogen bonds observed in the design model, likely due to slight differences in orientation of the binder to DNA and deviations from ideal B-DNA in the co-crystal structure. D43 was instead involved in a water-mediated hydrogen bond to C11 (**Fig. 3C**), and R49 was part of a hydrogen bond network involving the phosphate backbone. An additional water-mediated hydrogen bond was observed between S42 and A10. While water-mediated interactions are not considered by the Rosetta protocol used to build the side chains in the final design model, the LigandMPNN sequence design method implicitly considers these as the PDB training set contains many examples of water-mediated hydrogen bonds, which are known to confer additional specificity in native DBPs(*19*, *20*, *35*). Extensive hydrogen bond networks were also observed with the DNA phosphate backbone with most involved protein residues supported by sc-sc hydrogen bonds and packing interactions. These hydrogen bond networks with the phosphate backbone imply that much of the docking orientation is dominated by these interactions, suggesting that further enrichment for these features could improve design success rates.

**Fig. 3.**
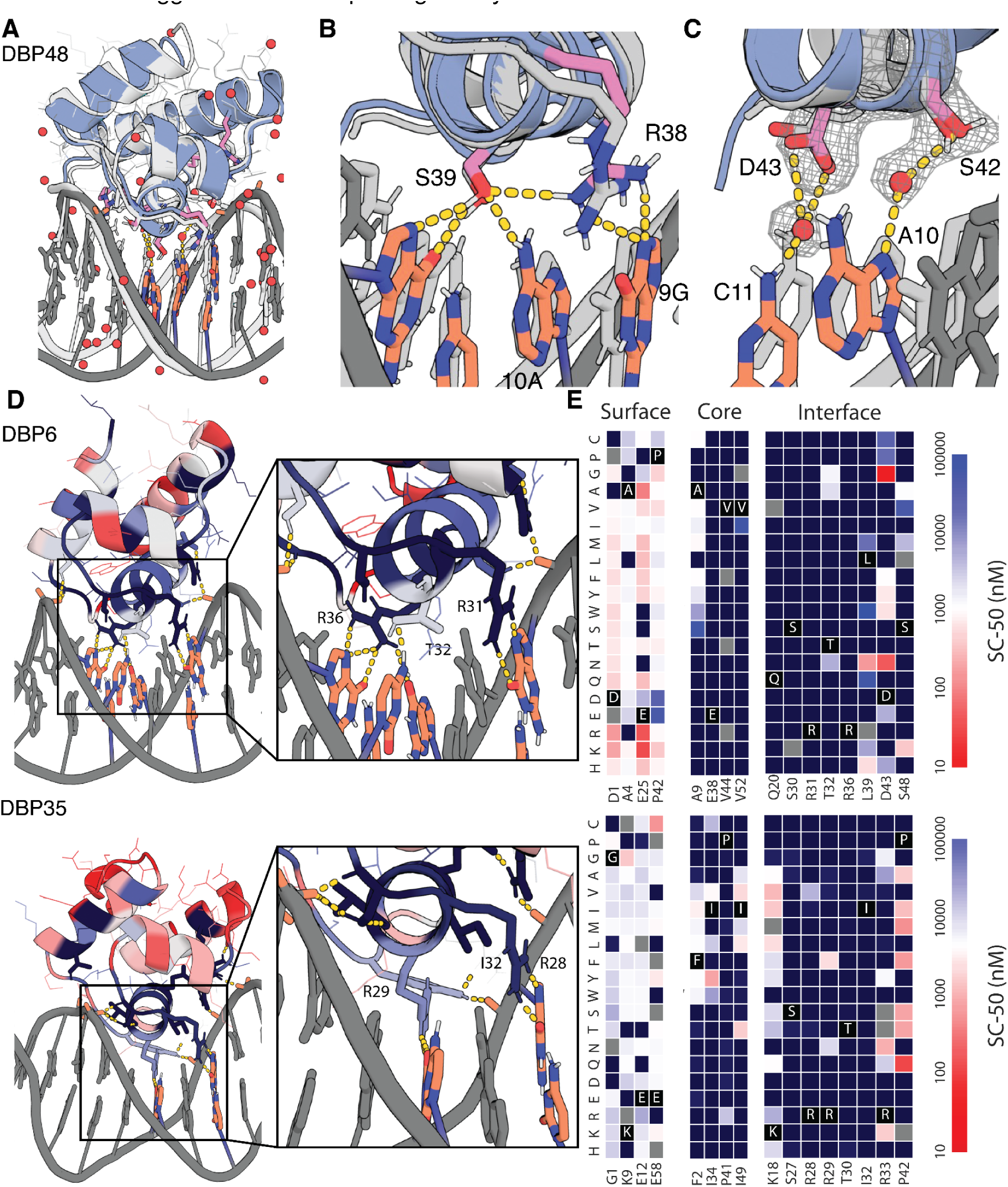
Structural validation of DNA binder designs. **(A)** Co-crystal structure of DBP48 (colored) and the design model (gray) are in close agreement. (**B**) Zoom-in showing the close agreement of critical interface residues R38 and S39 between the crystal structure and design model. (**C**) Close-up of water-mediated hydrogen bonds formed by S42 and D43. (**D**) (**Left**) Designed DNA binding proteins colored by positional Shannon entropy from site saturation mutagenesis, with blue indicating positions of low entropy (conserved) and red those of high entropy (not conserved). (**Right**) Zoomed-in views of central regions of the design interfaces. (**E**) Heat maps representing SC-50 values for single mutations in the design model core (left) and the designed interface (right). Substitutions that are heavily depleted are shown in blue, and beneficial mutations are shown in red. Full SSM maps over all positions and close-up views of DBP1 are provided in **fig. S12-13**.

To assess the contributions of each amino acid to binding for additional designs, we generated high-resolution footprints of the binding surface by sorting site saturation mutagenesis libraries (SSMs) in which every residue was substituted with each of the 20 amino acids one at a time for DBPs 1, 6, and 35 (**Fig. 3, D and E; fig. S12; and fig. S13**). For each of the three designs, we found that most positions at the interface and the core were largely conserved while positions at the surface were more tolerant of substitutions. In a small number of cases, substitutions led to notable improvements in binding affinity. The depletion of most substitutions in both the binding site and the core suggest that the design models are largely correct, whereas the enriched substitutions suggest routes to improving affinity.

### Assessment and optimization of designed DBP specificity

We selected 7 designs for further assessment of specificity *in vitro* using universal protein binding microarrays (uPBMs) containing all possible 7-mer DNA sequences(*36*, *37*). We found that several designs exhibited very high specificity for their design targets, most notably DBPs 6, 9, and 48 for which 7-mers containing the binding site motif were highly preferred over those lacking the motif (**Fig. 4A and fig. S14**). All-by-all analysis of on-target DBPs (**fig. S5**) showed that while some designs exhibit off-target binding, a number are highly specific to a single target. Five of our designed DBP-target pairs are highly orthogonal (**Fig. 4B and fig. S15i**).

**Fig. 4.**
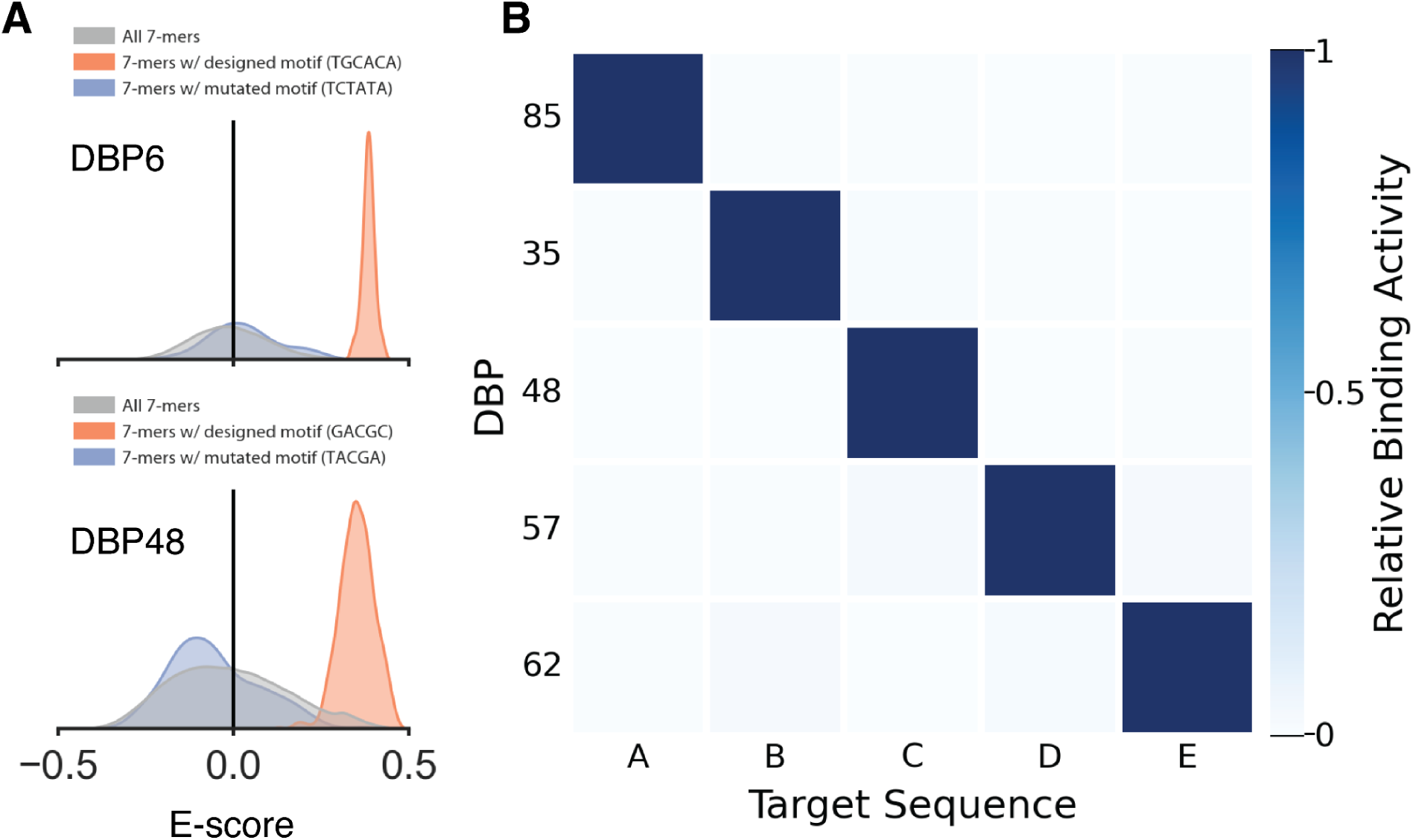
**Designed DBPs are highly specific.** **(A)** Histograms of E-score values for DBP6 (top) and DBP48 (bottom) from universal protein binding microarray experiments (uPBM) showing high specificity to the designed target sequence. The E-score distribution of all 7-mers are shown in gray while the distribution of 7-mers containing the designed binding site motif is shown in orange and the distribution for a mutated binding site motif is shown in blue. **(B)** All-by-all orthogonality matrix for 5 designed DNA binders screened by yeast display, normalized by row, at a DNA concentration of 1 µM (with avidity). Full orthogonality matrix with all tested DNA targets shown in **fig. S15I**.

For DBP35, which had moderate preference for the designed motif (**fig. S14**), we explored optimization of the specificity and affinity by combining substitutions found in mutational scanning. We found that combining three mutations (**fig. S15, A to C**) – R33N, which prevents a potential off target interaction; K18V, which adds an additional hydrophobic interaction with the methyl stem of base ADE11; and P42Q, which potentially stabilizes the protein scaffold structure – dramatically increased binding strength observed by yeast display with detectable binding down to ∼150 pM (**fig. S15D**). These mutations also increased specificity to 7 base positions as observed in a yeast competition assay (**fig. S15E**) compared with the 3 base position specificity observed in the original design (**Fig. 2B**), and substantially reduced binding to alternate motifs that were bound strongly in the original design (**fig. S1, F to H;** we refer to this optimized version as DBP35opt in the remainder of the text). Thus, just a few mutations of an initial design can lead to dramatic improvements in specificity and affinity.

### Designed DBPs modulate transcription in living cells

We tested the ability of our designed DBPs to function in cells to regulate transcription. To assay transcriptional repression in *Escherichia coli*, we constructed candidate NOT gates(*38*), where the input is a designed DBP under control of the IPTG-inducible P_Tac_ promoter and the output is yellow fluorescent protein (YFP) expression driven by a promoter incorporating the DBP DNA binding site (**fig. S16A)**. Single DBP domains and two copies of the same DBPs tethered through a flexible linker failed to exhibit YFP repression upon IPTG induction (**fig. S16B**), suggesting a need for higher affinity binding, longer sequence recognition, and/or a bulkier binding protein for effective hindrance of transcription initiation by *E. coli* RNA polymerase. To increase avidity and bulk, we positioned two copies of the same DBP (or one copy each of two different DBPs) on B-form DNA containing two palindromic copies of the target site (or the two different target sites), separated by different numbers of bases. We then used RFdiffusion(*39*) to build out new protein backbone segments that either transition into the TetR homodimer(*40*) or interact directly in homo- or heterodimeric arrangements (**Fig. 5A**). Following sequence design with ProteinMPNN to favor folding of the extensions to the intended structure and assembly of the intended dimers, we used AF2 to predict the structures of the homo- and heterodimers and selected those that were close to the design models. We experimentally characterized the ability of these designs to repress transcription from synthetic promoters incorporating two dimer binding sites (4 individual domain binding sites in total) flanking the −35 promoter region. Dose-dependent repression (> 2 fold) was observed for 2 of 96 TetR incorporating homodimeric designs (**fig. S16, C to E**) and 18 of 192 entirely *de novo* homodimeric and heterodimeric repressors incorporating different designed DBPs (**Fig. 5A and fig. S16, F and G**). All-by-all characterization of 6 selected designs and the corresponding cognate promoters showed considerable orthogonality (**Fig. 5, B and C**), with up to 20-fold repression for cells with the cognate target (DBP57_A2/DBP57_A2). In particular, two *de novo* dimeric designs with two copies of DBP57 designed to bind palindromic arrangements of the cognate target site at different spacings and in different orientations were each specific for their intended target, indicating that a single domain can serve as the basis for creating an array of orthogonal repressors.

**Fig. 5.**
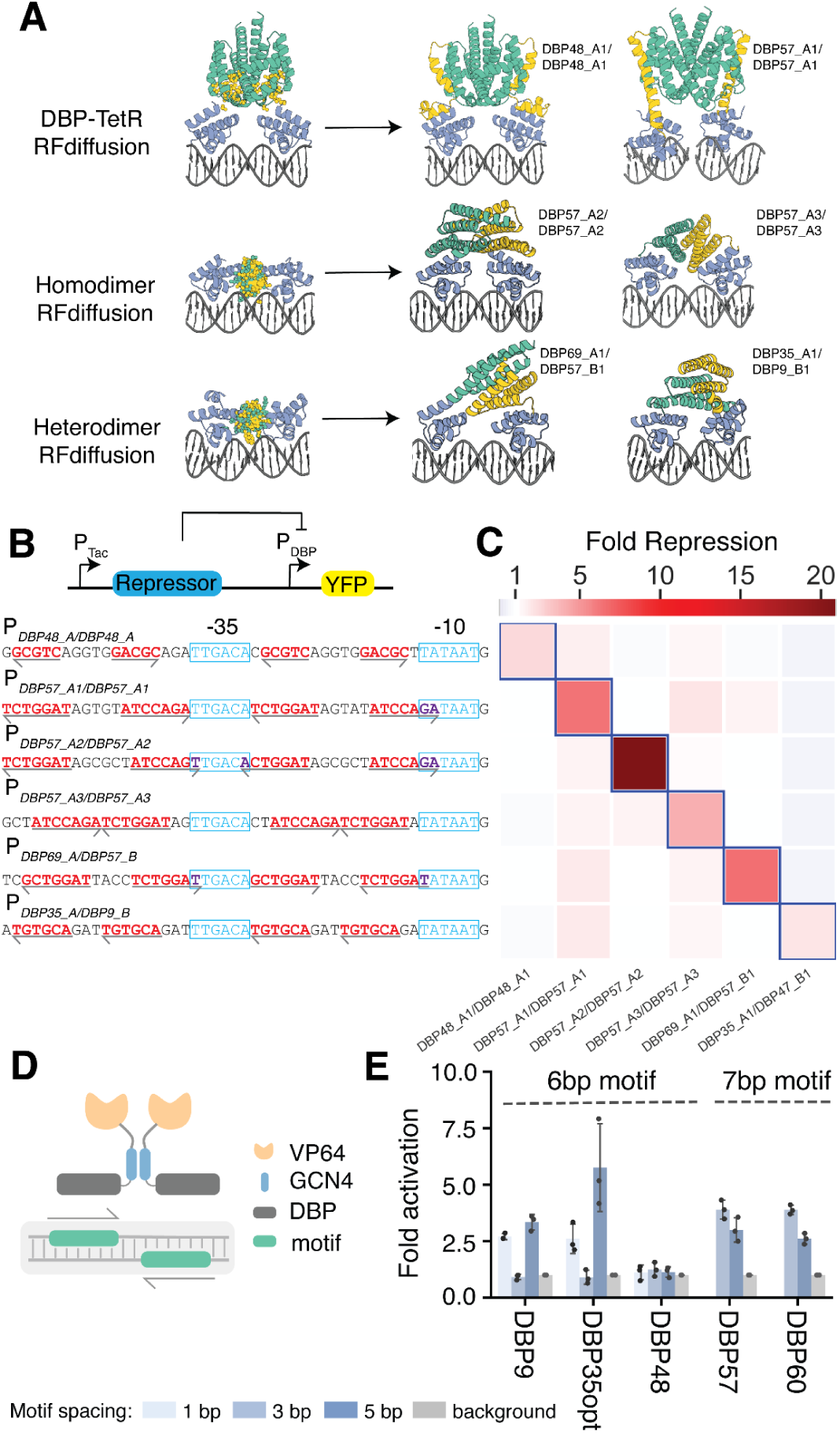
**Designed DBPs function in living cells to direct transcriptional repression and activation.** **(A)** Illustration of the RFdiffusion method for building out DBP domains into homo- or heterodimer arrangements, along with repressor designs selected for all-by-all repression assays. DBP48_A1/DBP48_A1 and DBP57_A1/DBP57_A1 are homodimer constructs transitioned into the TetR backbone, the remainder are *de novo* homo- or heterodimer constructs. (**B**) Transcriptional repression in *Escherichia coli.* Functional IPTG-inducible repressor block transcription of YFP from a synthetic promoter containing the designed DBP binding sites (red text) around the −10 and −35 elements (blue text). Arrows indicate directionality of the binding site. (**C**) All-by-all orthogonality matrix showing fold repression of YFP Fluorescence from flow cytometry analysis of cells containing the successful NOT gate circuits. Blue outlines indicate on-target repressor-promoter pairs. (**D**) Transcriptional activation in HEK293T cells measured by ENGRAM. synTFs were created by fusing the GCN4 dimerization domain and the VP64 activation domain to the C-termini of the DBPs. The synTF-specific cis-regulatory elements (CRE) were created by evenly distributing palindromic binding motifs on a 130 bp transcriptionally inactive DNA sequence where each CRE drives a uniquely barcoded pegRNA for recording into DNA TAPE. (**E**) Fold activation of synTFs measured as normalized barcode abundance.

Next, we tested DBP function as activators in mammalian cells. A set of synthetic transcription factors (synTFs) were created by fusing the GCN4 dimerization domain and the VP64 activation domain to the C-termini of DBPs 9, 35opt, 48, 57, and 60, which collectively recognize 3 unique motifs. The dimerization domain allows the DBPs to recognize a palindromic target sequence consisting of two binding motifs, increasing the binding affinity to the DNA sequence. We used the ENGRAM(*41*) recording technology to measure the activity of specific cis-regulatory elements (CREs) in HEK293 cells (**Fig. 5D**). In ENGRAM, each CRE drives expression of a uniquely barcoded pegRNA, which upon expression is recorded into the DNA TAPE at the HEK3 locus by prime-editor PEMax. After analyzing the barcode abundance for each individual CRE, we observed 3-5 fold activation for DBPs 9, 35opt, 57, and 60 (**Fig. 5E**).

### Determinants of DBP design success

Across all targets, designs that bound specifically to their intended target (**Fig. S4**) tended to have more sidechain- and mainchain-phosphate hydrogen bonds, lower Rosetta ΔΔG, and lower Cα RMSD of the AF2-predicted structure to the design model (**fig. S17**), while nonspecific binding was strongly correlated with a positive net charge. We did not observe enrichment of higher RotamerBoltzmann probabilities for base-hydrogen bonding sidechains, likely due to prior enrichment in the ordered design sets. However, we did observe enrichment of higher RotamerBoltzmann probabilities for phosphate hydrogen-bonding sidechains (**fig. S17**). Further enrichment for these metrics should increase design success rates.

A key feature of our design method is sampling from numerous diverse starting structures and docking positions to find complexes that can engage both the bases for sequence-specific recognition and the phosphate backbone to favor the designed binding mode. Like the most specific of our designs, native DNA binding proteins also have geometries enabling formation of mainchain-phosphate hydrogen bonds (**fig. S18**) and highly preorganized sidechain-phosphate hydrogen bonds (**fig. S2**). This is perhaps due to the inherent rigidity of these interactions which favor specific docks and restrict otherwise possible interactions of flexible sidechains with off-target DNA base atoms. To explore the importance of phosphate contacts mediating specific docks for achieving specificity to a given target site, we performed LigandMPNN redesign of 14 hits from our design campaigns against 100 randomly generated target sequences. Upon Rosetta relaxation of the redesigned complexes (20 LigandMPNN designed proteins per target-scaffold pair) in the presence of DNA, we observed that only 2 of the 100 sequences have as favorable Rosetta ΔΔGs and as many hydrogen bonds to bases (**fig. S19**), suggesting that the details of the scaffold backbone and dock make important indirect contributions to specificity by locking in the exact binding mode and narrowing the range of possible sidechain-base contacts. This makes it difficult to design DBPs to new DNA sequences through a native redesign approach.

## Conclusion

Our de novo DNA binder design approach can generate DBPs that specifically bind arbitrary DNA sequences, including sequences that are not bound by known DBPs in the PDB or JASPAR databases. These designed DBPs function both *in vitro* and in living cells, as observed through transcriptional repression and activation assays in both *Escherichia coli* and eukaryotic cells, respectively. The method samples structurally diverse HTH scaffolds to identify complexes that can facilitate specific contacts with the target DNA bases. The best designs were highly specific for their intended targets, and the crystal structure and specificity profiling assays strongly corroborate the computational design models.

The design method presented in this paper now enables the design of custom DNA binding miniproteins to target specific DNA sequences for diverse applications in gene regulation and editing. The ability to incorporate designed DBPs into transcriptional regulators through homo and hetero dimerization (**Fig 5**) should allow expansion of orthogonal TF-operator pairs for more complex gene circuits(*42*). TF designs composed of the same DBP domain orthogonally regulate distinct promoter sequences differing only in the spacing and orientation of the incorporated DNA binding sites, demonstrating the ability of RFdiffusion to precisely position two binding domains relative to one another. With RFdiffusion, it should be straightforward to fuse DNA-binding miniproteins together in a single chain in defined spatial orientations to allow specific targeting of longer target sites, or link DBPs with epigenetic modifiers or other effector recruiting domains to provide functionality beyond transcriptional activation and repression. Computationally designed DBPs are also well-suited for the simultaneous recognition of both DNA sequence and shape, including non-B DNA structures that may occur in ∼13% of the human genome(*43*). For any given backbone, there will likely be limitations in the ability to target certain DNA sequences, which likely explains the challenges in generating Cys_2_His_2_ zinc finger and other native scaffolds to bind to some target sites; our approach provides a powerful new method for building DNA binding proteins with backbones tailored to specific sequences of interest, and we anticipate the method and the sequence specific designs should be widely useful in synthetic biology and other areas requiring sequence specific DNA recognition.

## Methods

### Scaffold Library Generation

Scaffolds deposited in the PDB with structural similarity to selected template backbones (PDB IDs: 1L3L(*44*), 1PER(*45*), 1EFA(*46*), 1DDN(*47*), and 1APL(*48*)) were identified using TM-align(*28*). Amino acid sequences of identified protein scaffolds were used as seeds to generate multiple sequence alignments (MSAs) using an HHBlits(*49*) search of the UniRef30 database(*50*). Resulting MSAs were used for HMMer(*51*) searches of the JGI metagenome protein sequence databases(*52*) and the Uniref100 database(*50*). HMMer search results were clustered to < 70% sequence identity using MMSeqs2(*53*) and MSAs were generated from each clustered sequence using HHBlits. AlphaFold2(*27*) was used to predict structures for each sequence using the generated MSAs. Resulting scaffolds were filtered for high confidence AlphaFold2 pLDDT scores, TMscore to the input backbone templates, and Rosetta score. Scaffolds of specific topologies were supplemented with additional AlphaFold2-predicted structures of transcription factor sequences identified from bacterial metagenomes using DeepTF(*54*). PSSMs were generated for each scaffold using PSI-Blast(*55*) and custom code for use as constraints of Rosetta design. All final scaffolds are available for download.

### RIF docking of scaffolds onto DNA targets

Structures of B-DNA were generated by either (1) using the DNA portion of PDB structures 1BC8(*56*), 1YO5(*57*), 1L3L(*44*), 2O4A(*58*), 1OCT(*59*), 1A1F(*60*), and 1JJ6(*61*), or (2) using the software X3DNA(*62*), followed by a constrained Rosetta relax of the DNA structure. The RIF docking method performs a high-resolution search of continuous rigid-body docking space. RIF docking comprises two steps. In the first step, ensembles of interacting discrete sidechains (referred to as ‘rotamers’) tailored to the target are generated. Polar rotamers are placed on the basis of hydrogen-bond geometry whereas apolar rotamers are generated via a docking process and filtered by an energy threshold. Rotamers were only calculated for nucleotide base atoms in the major groove of the DNA target. All the RIF rotamers are stored in ∼0.5 Å sparse binning of the six-dimensional rigid body space of their backbones, allowing extremely rapid lookup of rotamers that align with a given scaffold position. To enrich for canonical protein-DNA hydrogen bond interactions, rotamers of ARG, GLN, and ASN forming bidentate hydrogen bonds with G and A bases were extracted from the PDB, clustered by RMSD, aligned to the DNA target at all G and A positions, and added to the RIF as hotspot residues. To facilitate the next docking step, RIF rotamers are further binned at 1.0 Å, 2.0 Å, 4.0 Å, 8.0 Å and 16.0 Å resolution. In the second step, a set of scaffolds is docked into the produced rotamer ensembles, using a hierarchical branch-and-bound search strategy. Starting with the coarsest 6.0 Å resolution, an enumerative search of scaffold positions is performed: the designable scaffold backbone positions are checked against the RIF to determine whether rotamers can be placed with favorable interacting scores. All acceptable scaffold positions (up to a configurable limit, typically ten million) are ranked and promoted to the next search stage. Each promoted scaffold is split into 26 child positions in the six-dimensional rigid-body space, providing a finer sampling. The search is iterated at 8.0 Å, 4.0 Å, 2.0 Å, 1.0 Å and 0.5 Å resolutions. All RIF docks were required to utilize at least 1 hotspot residue to be saved as an output.

### Energy function optimization

A new version of the Rosetta score function was trained to better evaluate the energy of protein-DNA interfaces. Additional flexibility of the DNA duplex was incorporated into Rosetta’s rotamer optimization and gradient-based minimization modules using modifications of DNA dihedral angles(*63*) and the score function was optimized using the same general method as previously published(*64*). The weights of individual terms in the score function were optimized to reproduce the geometries of DNA crystal structures. Specifically, the distributions of pairwise atomic distances, base-stacking and base-pairing geometries, and bond torsions were considered. Additional optimization was performed on tasks related to protein-DNA complex structures. These tasks included energy ranking of perturbed crystal structures, rotamer recovery in repacking crystal structures, and sequence recovery in redesigning the protein sequence of crystal structures. An additional weight was placed on the frequency of positively charged residues at interface positions, because previous score functions tended to overestimate the strength of solvent-exposed charged interactions. Similar geometric and design tasks were included for protein structures alone. Rosetta score weights optimized included partial atomic charges of protein and DNA, hydrogen bond strengths, and solvation energies. The resulting score function showed improvement across nearly all tasks, with the greatest improvements found in the protein-DNA energy ranking and sequence design.

### RotamerBoltzmann filters

The Boltzmann probability of finding a given rotamer in a specific state was evaluated using the RotamerBoltzmannWeight filter in Rosetta(*30*). The RotamerBoltzmann score is an approximation of preorganization of a given residue in the unbound state. All amino acid residues forming hydrogen bonds with DNA base or phosphate atoms were evaluated by this metric, which was calculated on the protein monomer in the unbound state. The metric was estimated by fixing neighboring sidechains and assessing the Boltzmann probability distribution on rotamers accessible by the sidechain of interest. In order to increase the likelihood of a given rotamer in the protein-DNA complex, designs with lower RotamerBoltzmann scores (a score of 0 implies the rotameric state is unpopulated and a score of 1 implies the state is the only populated state) were preferentially chosen, as known native protein-DNA crystal structures tend to contain preorganized amino acid residues (**fig. S2**).

### Rosetta-based interface sequence design

A stripped down version of the Rosetta score function was used to roughly design the interface of RIF dock outputs(*5*). This step was primarily used to replace clashing residues before evaluating for design potential. Specifically, fa_elec, lk_ball[iso,bridge,bridge_unclp], and the _intra_ terms were disabled. All that remained were Lennard-Jones, implicit solvation and backbone-dependent one-body energies (fa_dun, p_aa_pp, rama_prepro). Additionally, flags were used to limit the number of rotamers built at each position (Supplementary Information). After the rapid design step, the designs were minimized twice: once with a low-repulsive score function and again with a normal-repulsive score function. Rosetta ΔΔG and contact molecular surface were then calculated on the roughly designed interface. A maximum likelihood estimator was used to give each predicted design a likelihood that it should be selected to move forward. A subset of the docks to be evaluated were subjected to the full sequence design, and their final metric values calculated. With a goal threshold for each filter, each fully designed output can be marked as pass or fail for each metric independently. Then, by binning the fully designed outputs by their values from the rapid trajectory and plotting the fraction of designs that pass the goal threshold, the probability that each predicted design passes each filter can be calculated. From here, the probability of passing each filter may be multiplied together to arrive at the final probability of passing all filters. This final probability can then be used to rank the designs and pick the best designs to move forward to full sequence optimization. Note that the rapid design protocol here is used merely to rank the designs, not to optimize them; the original docks are the structures carried forward.

These docked conformations passing the rapid design protocol were further optimized to generate shape- and chemically-complementary interfaces using a Rosetta FastDesign protocol, alternating between sidechain rotamer optimization and gradient descent–based energy minimization. Design was performed with a sequence profile constraint based on an MSA of the originating native scaffold sequence and cross-interface interactions upweighted to maximize contacts and shape complementarity. We did not allow Rosetta to repack or relax the DNA target during the design procedure. A python script was implemented to automatically carry out rapid design evaluation, pre-emption, and full sequence design. Computational metrics of the final design models were calculated using Rosetta, which includes ΔΔG, hydrogen bonds to base atoms, and contact molecular surface, among others, for design selection. All the script and flag files to run the programs are provided in the Supplementary Information. ProteinMPNN was used to redesign non-interface residues in the final design step, before AF2 monomer validation.

### LigandMPNN-based sequence design

LigandMPNN was used for sequence design in the context of DNA. The network was used to optimize the protein sequence for given protein-DNA complex structures during design, whereby amino acids were determined autoregressively by the identity and location of neighboring protein and DNA residues. When the full protein sequence was determined, it was threaded onto the input protein scaffold. As in the above Rosetta-based interface sequence design protocol, the designs were minimized with a low-repulsive score function and again with a normal-repulsive score function, and Rosetta ΔΔG and contact molecular surface were calculated on the roughly designed interface. A maximum likelihood estimator was used to pre-empt design of poor docks as described in the above Rosetta-based sequence design protocol. A python script was implemented to automatically carry out MPNN sequence design, rapid design evaluation, pre-emption, and Rosetta Relax. Computational metrics of the final design models were calculated using Rosetta, which includes ΔΔG, interface hydrogen bonds, and contact molecular surface, among others. LigandMPNN temperatures of 0.2–0.3 were used earlier in the design process to increase the variability of amino acid sequences, while a temperature of 0.1 was used later to determine the more probable sequences. Key residues making base-specific hydrogen bonds with DNA atoms were fixed in later stages of the pipeline to encourage the design of supporting residues. All the script and flag files to run the programs are provided in the Supplementary Information.

### Backbone resampling with motif grafting

Motif grafting was performed as previously reported(*5*). Briefly, the binding energy and interface metrics for all the continuous secondary structure motifs (helix, strand and loop) were calculated for the designs generated in the broad search stage, as performed in previous work(*5*). The motifs with good interactions (based on binding energy and other interface metrics, such as contact molecular surface) with the target were extracted and aligned using the target structure as the reference. All the motifs were then clustered based on an energy-based TM-align-like clustering algorithm(*28*) without any further superimposition. The best motif from each cluster was then selected based on the per-position weighted Rosetta binding energy, using the average energy across all the aligned motifs at each position as the weight. Around 500–2,000 best motifs were selected, and the scaffold library was superimposed onto these motifs using the MotifGraft mover(*65*). Interface sequences were further optimized, and computational metrics were computed for the final optimized designs as described in the Rosetta- and LigandMPNN-based sequence design methods.

### Backbone remodeling with protein inpainting

Scaffold secondary structures were determined using DSSP(*66*). ProteinInpainting contigs were generated for each design that mask scaffold loops longer than 4 residues and surrounding residues, while ensuring that all residues forming hydrogen bonds to the DNA backbone were conserved. 10–20 unique contigs were generated for each design and sequences were constrained to a maximum of 65 amino acids. ProteinInpainting outputs were aligned to the DNA target using fixed interface residues of the input structure. The aligned ProteinInpainting outputs were subject to several further LigandMPNN + FastRelax rounds before AF2 monomer prediction and superposition steps.

### AF2 monomer validation and superposition

AF2 structures were produced using the single sequence of each design. AF2 was run with model 1 and 12 recycles for each design. C-alpha RMSD of the AF2 structures to each respective design model were calculated. AF2 structures were superpositioned onto the DNA target using the backbone coordinates of interface residues within 8 Å of the DNA target. A fixed backbone Rosetta FastRelax was performed on each superpositioned complex and all relevant metrics were calculated on the final superpositioned design model.

### Design Filtering

Designs were filtered after each sequence design step and after superimposition of AlphaFold2 models for those with the most favorable free energy of binding (Rosetta ΔΔG), contact molecular surface area(*5*) and interface hydrogen bonds, the fewest interface buried unsatisfied hydrogen bond donors and acceptors, and those containing bidentate sidechain-base hydrogen bonding arrangements frequent in the PDB, including bidentate interactions of ARG-G, GLN-A, and ASN-A. Designs were additionally filtered for those with a high RotamerBoltzmann score among ARG, LYS, GLN, or ASN residues forming hydrogen bonds with bases (max rboltz RKQE) and those with a high median RotamerBoltzmann (median rboltz) score of all residues forming hydrogen bonds with bases.

### DNA library preparation

All protein sequences were padded to 65 amino acids by adding a (GGS) n linker at the carboxy terminus of the designs to avoid the biased amplification of short DNA fragments during PCR reactions. The protein sequences were reversed translated and optimized using DNAworks2.0(*67*) with the Saccharomyces cerevisiae codon frequency table. Oligonucleotide pools encoding the designs were purchased from Agilent Technologies.

All libraries were amplified using Kapa HiFi polymerase (Kapa Biosystems) with a qPCR machine (Bio-Rad, CFX96). In detail, the libraries were first amplified in a 25 μl reaction, and the PCR reaction was terminated when the reaction reached half maximum yield to avoid overamplification. The PCR product was loaded onto a DNA agarose gel. The band with the expected size was cut out, and DNA fragments were extracted using QIAquick kits (Qiagen). Then, the DNA product was re-amplified as before to generate enough DNA for yeast transformation. The final PCR product was cleaned up with a QIAquick Clean up kit (Qiagen). For the yeast transformation step, 2–3 µg of linearized modified pETcon vector (pETcon3) and 6 µg of insert were transformed into the EBY100 yeast strain using a previously described protocol(*68*).

DNA libraries for deep sequencing were prepared using the same PCR protocol, except the first step started from yeast plasmid prepared from 5 × 10^7^ to 1 × 10^8^ cells by Zymoprep (Zymo Research). Illumina adapters and 6-bp pool-specific barcodes were added in the second qPCR step. Gel extraction was used to obtain the final DNA product for sequencing. All the different sorting pools were sequenced using Illumina NextSeq sequencing.

### Yeast surface display

*Saccharomyces cerevisiae* EBY100 strain cultures were grown in C-Trp-Ura medium supplemented with 2% (w/v) glucose. For induction of expression, yeast cells were centrifuged at 6,000*g* for 1 min and resuspended in SGCAA medium supplemented with 0.2% (w/v) glucose at the cell density of 1 × 107 cells per ml and induced at 30 °C for 16–24 h. Cells were washed with PBSF (PBS with 1% (w/v) BSA) and labeled with biotinylated targets using two labeling methods: with-avidity and without-avidity labeling. For the with-avidity method, the cells were incubated with biotinylated target, together with anti-c-Myc fluorescein isothiocyanate (FITC, Miltenyi Biotech) and streptavidin–phycoerythrin (SAPE, ThermoFisher). The concentration of SAPE in the with-avidity method was used at one-quarter of the concentration of the biotinylated targets. For the without-avidity method, the cells were first incubated with biotinylated targets, washed and secondarily labeled with SAPE and FITC.

Cell sorting of labeled yeast pools was performed using a Sony SH800S cell sorter. Libraries of designs were sorted using the with-avidity method for the first few rounds of screening to exclude weak binder candidates, followed by several without-avidity sorts with different concentrations of targets. For SSM libraries, two rounds of with-avidity sorts were applied and in the third round of screening the libraries were titrated with a series of decreasing concentrations of targets to enrich mutants with beneficial mutations.

For yeast display characterization of individual designs, including competition assays, DNA sequences encoding the proteins of interest were purchased as Integrated DNA Technologies (IDT) E-Blocks, transformed into yeast cells, and incubated in 96 well culture plates. Labeling with biotinylated dsDNA targets and SAPE/FITC was performed in a 96 well plate format. Of the 44 designs which were confirmed to bind their intended target in clonal yeast display experiments, (**fig. S4**), we categorized 14 with detectable binding to less than 3 of the 13 tested DNA targets (**fig. S5**) as specific binders and the remainder as nonspecific.

For yeast display competition assays, labeling was performed without avidity using 1 µM biotinylated dsDNA duplex oligos and an excess of 8 µM non-biotinylated competitor dsDNA duplex oligos. As indicated in figure captions, some competition assays for higher affinity binders were carried out with lower dsDNA oligo concentrations. Flow cytometry analysis was performed with an Attune NxT flow cytometer with autosampler. Flow cytometry data analysis was performed using custom python code and the CytoFlow python package. For each individual sample, gating of the expression population was performed using the CytoFlow Gaussian Mixture Model and the ratio of SAPE channel intensity to FITC channel intensity (binding signal / expression signal) was calculated for all gated expression events of the sample.

### Deep sequencing analysis

The Pear program was used to assemble the fastq files from the deep sequencing runs. Translated, assembled reads were matched against the ordered design to determine the number of counts for each design in each pool. In each sequenced pool, binder enrichment was calculated by determining the percent of reads for each binder design in the pool and dividing this number by the same value in the naive expression sort pool. Designs were considered binders if > 100-fold enrichment was observed in the last 1 µM with-avidity sort to the designed dsDNA target. For SSM libraries, apparent SC50 was estimated using the fitting procedure described in Longxing et al.(*5*)

### Protein expression and purification

DNA sequences encoding the proteins of interest were purchased as Integrated DNA Technologies (IDT) E-Blocks and incorporated into plasmids using Golden Gate assembly. The plasmids were then transformed into BL21(DE3) competent *E. coli*. The transformation reactions were used to inoculate starter cultures in 5 mL or 25 mL of “Terrific Broth” (TB), supplemented with 1% (w/v) glucose and 50 mg/L kanamycin. After shaking overnight at 37°C, the starter cultures were diluted 50-fold into 50 mL or 500 mL of TB with kanamycin. These cultures were incubated at 37°C, shaking, until the optical density (OD) reached 0.6-0.8, at which point protein expression was induced by the addition of IPTG. The cultures were then further incubated overnight at 18°C. Cells were harvested by centrifugation for 15 min at 3000*g*, pellets resuspended in lysis buffer (150 mM NaCl, 20 mM Tris-HCl, 0.5 mg/mL DNAse I, 1 mM PMSF, pH 8.0), the cells lysed by sonication, and the lysate clarified by further centrifugation for 30 min at 20,000*g*. The supernatant was passed through Ni-NTA resin in a gravity column, and then the resin was washed with 20 column volumes of high-salt wash buffer (2 M NaCl, 20 mM Tris-HCl, 20 mM Imidazole, pH 8.0). Either (A) the His-tagged protein was eluted with 2 column volumes of elution buffer (1 M NaCl, 20 mM Tris, 250 mM Imidazole, pH 8.0), or (B) the resin was further washed with 5 column volumes of SNAC buffer (100 mM CHES, 100 mM Acetone oxime, 100 mM NaCl, 500 mM GnCl, pH 8.6), incubated in 5 column volumes of SNAC buffer + 0.2 mM NiCl_2_ on an orbital shaker at room temperature overnight, and collected as the column flow-through. Whether cleaved or not, the protein was concentrated to about 1 mL and loaded in 500 μL samples onto a Cytiva Superdex™ 75 Increase 10/300 GL gel filtration column equilibrated in buffer (1 M NaCl, 20 mM Tris-HCl, pH 8.0). Fractions containing monomeric protein were pooled and concentrated to about 200 μL. Protein concentrations were estimated spectroscopically by absorbance at 280 nm. For proteins with no Trp, Tyr, or Cys residues, concentrations were approximated by Bradford reagent absorbance at 470 nm in comparison to BSA standards of known concentration.

### Biolayer interferometry

Biolayer interferometry binding data were collected on an Octet R8 (Sartorius) and processed using the instrument’s integrated software. Biotinylated dsDNA oligos were loaded onto streptavidin-coated biosensors (ForteBio) at 200 nM in PBS + 1% BSA + 0.05% Tween 20 for 6 min. Analyte proteins were diluted from concentrated stocks into the binding buffer. After baseline measurement in the binding buffer alone, the binding kinetics were monitored by dipping the biosensors in wells containing the target protein at the indicated concentration (association step) and then dipping the sensors back into baseline/buffer (dissociation). Data were analyzed and processed using ForteBio Data Analysis software v.9.0.0.14.

### Crystallization and Structure Determination

Purified DBP48 was complexed with duplex DNAs, of varying duplex length and a single 5’overhang base, to a final concentration of 176 µM DBP48 and 233 µM duplex DNA. Complexes were screened for crystals in several broad matrix screens using a mosquito robot (SPT LabTech) then possible hits were optimized in 24-well hanging drop trays with a 2 micro-liter drop containing a 1:1 ratio of complex to well solution and equilibrated over 1 mL of well solution. A single diffraction quality crystal was obtained with duplex DNA of length 10 basepairs (5’-ACCTGACGCGA-3’, 3’-GGACTGCGCTT-5’) and a well condition containing 200 mM ammonium acetate, 100 mM sodium acetate at pH 4.6, and 28% Polyethylene glycol 4000. The crystal was washed in well solution then flash frozen directly by plunging into liquid nitrogen. Data was collected at the Advanced Light Source in Berkeley, CA on beam line 5.0.1 at a wavelength of 0.9762 Å and processed with DIALS(*69*). Phases were determined via molecular replacement by searches with the original computational protein design and duplex DNA using Phaser(*70*) in the Phenix suite(*71*). The top scoring molecular replacement solutions were run through a round of refinement with Phenix refine and further rounds of refinement with Phenix refine and rebuilding with Coot(*72*) were performed on the top scoring structure. Data collection and refinement statistics are reported in **table S4**.

### Universal protein binding microarrays (uPBMs)

Universal PBM experiments were carried out following the standard PBM protocol(*36*, *37*). Briefly, we first performed primer extension to obtain double-stranded DNA oligonucleotides on the microarray. Next, each microarray chamber was incubated with a 2% milk blocking solution for 1 h, followed by incubations with a PBS-based protein binding mixture for 1 h and with Alexa488-conjugated anti-His antibody (1:20 dilution, Qiagen 35310) for 1 h. The array was gently washed as previously described(*36*) and then scanned using a GenePix 4400A scanner (Molecular Devices) at 5-μm resolution. Data were normalized and processed with standard analysis scripts(*36*, *37*).

### RFdiffusion-based design of DBP-TetR fusion linkers, homodimers, and heterodimers

For TetR fusions, diffusion inputs were generated by manually aligning DBP domains (DBPs 48, 57, and 69) symmetrically relative to the TetR homodimer scaffold. 10,000 RFdiffusion trajectories were run per input to generate rigid linkers between the DBP domains and the TetR homodimer scaffold. ProteinMPNN sequence design was performed on dimer diffusion outputs with tied positions between the two units and most residues of the DBP fixed, only allowing design of DBP residues nearby the newly diffused linker region. Homodimer complexes were predicted with ESMFold due to the inability of AF2 to predict the MPNN-designed TetR backbones. Predicted structures were filtered on RMSD of the predicted DBP regions to the input DBP domains and ESMFold pLDDT to select 96 designs across the three inputs. For homodimer and heterodimer design, diffusion inputs were generated by aligning DBP domains (DBPs 9, 35opt, 57, and 69) symmetrically or asymmetrically onto DNA. 10,000 RFdiffusion trajectories were run per input to generate C2-symmetric homodimers or asymmetric heterodimers between the DBP domains. ProteinMPNN sequence design was performed on diffusion outputs with tied positions between the two units (for homodimers) and most residues of the DBP fixed. Complexes were predicted with AlphaFold2 and filtered on RMSD of the predicted DBP regions to the input DBP domains and pLDDT to select 96 homodimer designs and 96 heterodimer designs.

### Transcriptional repression assays in *E. coli*

The pRF-TetR vector(*38*) was used for transcriptional repression assays in *E. coli.* A new version of this vector (pRF-BsmB1) was constructed by first removing the LuxR gene and then replacing the TetR gene, its terminator sequence, and regulated promoter with two BsmB1 cut sites such that new repressor variants and their associated promoters could be easily inserted via Golden Gate Assembly(*73*). For DBPs tethered with a flexible linker, a flexible linker was used to connect the C- and N- termini of two copies of the DBP (linker1: KESGSVSSEQLAQFRSLD, linker2: EGKSSGSGSESKST, linker3: GGGGGGGG, linker4: GSGSGSGSGSGSGSGS). Synthetic promoters were designed by inserting DNA binding sites around the consensus −10 and −35 elements of the *E. coli* RNAP promoter. Genes encoding the single domain DBP, flexibly linked, TetR fusions, homodimers, and heterodimers were ordered as Twist synthetic gene fragments encoding the repressor gene (using Twist codon optimization), a transcriptional terminator, and an associated synthetic promoter. Heterodimer constructs were encoded into bicistronic operons. Gene fragments were ordered containing BsmB1 cut sites on either end to allow for assembly into the modified pRF-BsmB1 vector. Upon Golden Gate assembly with the BsmB1 Type II-S restriction enzyme, plasmids were transformed into NEB 5-alpha competent *E. coli* cells and streaked onto Luria-Burtani (LB) plates containing carbenicillin. All-by-all repressor constructs (**Fig. 5c**) were cloned by digestion with BsiWI-HF (NEB) and BbsI (NEB), gel extraction of the backbone and promoter bands, followed by ligation with T4 DNA ligase and transformation into NEB 5-alpha competent *E. coli*.

Individual transformants were picked and verified via sanger sequencing. Sequence verified colonies were inoculated into 200 µL LB media containing carbenicillin for overnight growth in 96-well round bottom plates at 37℃ in a plate shaker. The following day, 2 µL of overnight cultures were transferred into a new plate containing 200 µL LB media containing carbenicillin and appropriate concentrations of Isopropyl ß-D-1-thiogalactopyranoside (IPTG) (1 mM in **Fig. 5c**) and grown for ∼ 18 hrs in 96-well round bottom plates at 37℃. Flow cytometry analysis of cultures was performed with an Attune NxT flow cytometer with autosampler. Flow cytometry data analysis was performed using custom python code and the CytoFlow python package. For each individual sample, gating was performed using the single component CytoFlow Gaussian Mixture Model and median BL1-A channel fluorescence was determined for all gated expression events of each sample. The median BL1-A channel fluorescence value of empty cells without a pRF vector was subtracted from the median BL1-A value of each sample. For each repressor variant in **Fig. 5c** and **fig. S16d**, fold repression was calculated from at least 7 biological replicates as the ratio of median BL1-A channel fluorescence of the uninduced sample (background subtracted) to the median BL1-A channel fluorescence of the induced sample (background subtracted).

### Transcriptional activation in HEK293T cells

HEK293T cells expressing the PEmax were cultured in DMEM High glucose (GIBCO), supplemented with 10% Fetal Bovine Serum (Rocky Mountain Biologicals) and 1% penicillin-streptomycin (GIBCO). Cells were grown with 5% CO_2_ at 37°C. 1 x 10^5^ cells were seeded on a 48-well plate a day before transfection. Enhancer plasmid and binder plasmid were mixed with a ratio of 2:1. Enhancer variants and background control were mixed with a ratio of 2:2:2:1. A total of 300 ng of plasmid were transfected using Lipofectamine 3000 (ThermoFisher, L3000015), following the manufacturer’s protocol. 3 synTF specific recorders and 1 TCF-LEF- recorder (negative control) were mixed with ratio 2:2:2:1 and co-transfected with synTFs into the HEK293T cells expressing PEmax. 3 different spacings were tested—1 bp, 3 bp, and 5 bp—between the palindromic binding motifs to maximize the recorder activity. Cells were harvested and analyzed 2 days post-transfection.Cells were harvested 2 days post-transfection. Genomic DNA was extracted based on the protocol described earlier(*41*). Briefly, cells were lysised using freshly prepared lysis buffer (10 mM Tris-HCl, pH 7.5; 0.05% SDS; 25 μg/ml protease (ThermoFisher)) for each well. The genomic DNA mixture was incubated at 50°C for 1 h, followed by an 80°C enzyme inactivation step for 30 min. The DNA TAPE was amplified from the genomic DNA directly for next generation sequencing. Recorded information was extracted via custom analysis code. Each enhancer has a unique barcode representing its activity. Transcription activation was measured as the fold change in the barcode abundance relative to the negative control barcode. All measurements were performed in triplicates. Error bars represent standard deviation of the mean relative barcode abundance.

## Data and materials availability

The underlying data and PDB files will be made available upon reasonable request to the authors. Scripts for RIFgen, RIFdock, and LigandMPNN sequence design will be made available upon reasonable request. The co-crystal structure of DBP48 has been deposited to RCSB as PDB ID 8TAC.

## Acknowledgements

We thank Nate Bennett for use of python scripts in the design pipeline, Justas Dauparas for LigandMPNN method development, and Kandise VanWormer for laboratory support. pRF-TetR was a gift from Christopher Voigt (Addgene plasmid # 49374; http://n2t.net/addgene:49374; RRID:Addgene_49374). HEK293T cells expressing PEmax was a gift from the Shendure lab.

## Author Contributions

C.G., R.P., R.M., D.C., F.D., and D.B. designed the research. C.G., R.P., and R.M. developed the computational binder design pipeline. R.M., H.H., D.C., and F.D. contributed to energy function optimization. C.G., R.P., I.G., and D.V. performed yeast screening, expression, and binding experiments. R.M. and C.G. performed *E. coli* protein expression experiments. C.G. performed biolayer interferometry experiments. O.B. performed universal protein binding microarray experiments. L.D. performed x-ray co-crystallography experiments. C.N. and G.L. developed the scaffold library generation method and the PSSM-based Rosetta constraints. R.P and C.G. implemented the LigandMPNN sequence design method for DNA binder design. B.C. contributed to computational methods development. C.G., B.L., E.N., and Y.P. designed repressors and performed *E. coli* transcriptional repression experiments. W. C. performed mammalian cell transcriptional activation experiments. All authors analyzed data. R.G., B.S., F.D., and D.B. supervised research. C.G., R.P., R.M., W.C., and D.B. wrote the manuscript with input from the other authors. All authors revised the manuscript.

## List of Supplementary Materials

**Fig. S1.** Overview of scaffold library generation.

**Fig. S2.** Native protein-DNA structures are enriched with highly preorganized residues forming hydrogen bonds with base atoms and the DNA phosphate backbone.

**Fig. S3.** Distribution of metrics calculated on each ordered design set.

**Fig. S4.** Clonal analysis of binder designs by yeast surface display confirms dsDNA-binding function.

**Fig. S5.** All-by-all analysis of selected designs by yeast surface display reveals preferential target binding of designs.

**Fig. S6.** Full competition assays for all DBPs designs. Fig. S7. SEC traces of purified proteins.

**Fig. S8.** Purified designs bind their respective dsDNA targets *in vitro* by biolayer interferometry. Fig. S9. Comparison of designed DBPs with nearest native structures by target motif or protein structure.

**Fig. S10.** Comparison of designed DBP target motifs to JASPAR database. Fig. S11. Global view of the DBP48 co-crystal structure.

**Fig. S12.** Full SSM maps for DBP1, DBP6, and DBP35. Fig. S13. Close-up view of DBP1 SSM.

**Fig. S14.** Analysis of DBPs 1, 3, 5, 6, 9, 48, and 35 with universal protein binding microarray experiments containing all 7-mers.

**Fig. S15.** Optimizing DBP35 to disrupt off-target DNA binding.

**Fig. S16.** Use of DBPs to direct transcriptional repression in *E. coli*.

**Fig. S17.** Power of computational metrics to predict binders.

**Fig. S18.** Mainchain-phosphate hydrogen bonds fix specificity and limit alternative target sites of DBP scaffolds.

**Fig. S19.** Starting dock restricts possible target sequences of DBP designs.

**Table S1.** Sequences of DNA binder designs, corresponding interface knockout positions, and aligned best hit of blastp search of the NCBI standard non-redundant protein database.

**Table S2.** Sequences used as dsDNA targets.

**Table S3.** PDB IDs for native DBPs similar to designs. Table S4. Data collection and refinement statistics.

