## Supplemental Information for "Computational design of sequence-specific DNA-binding proteins"

**Supplementary Note 1:**

Experimental characterization of a first round of designs generated using three-helix bundle scaffolds similar to those used for general protein binder design(*5*) showed that few bound their DNA targets and none bound specifically. To understand the reason for this failure, we carried out detailed structural inspection of these designs compared to natural DBPs. We observed that most natural DBPs make backbone amide-mediated hydrogen bond interactions with DNA phosphate oxygens (hereon called mainchain-phosphate hydrogen bonds) that were rare or absent in our docked and designed complexes. Thus, these scaffolds were unable to overcome the first DNA specific design challenge described in the design strategy.

**Figs. S1 to S19**

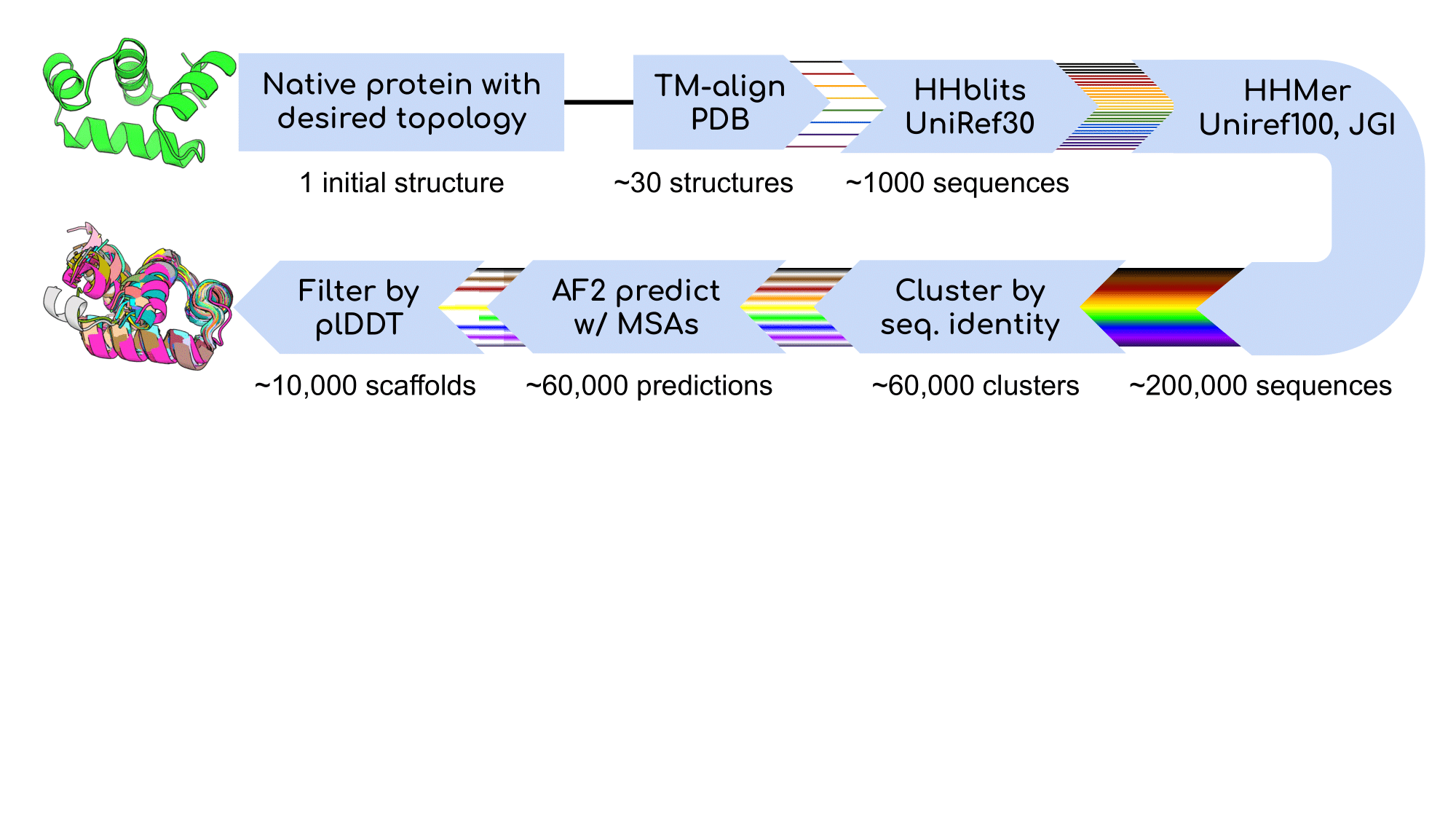

**Fig. S1. Overview of scaffold library generation.** Scaffolds deposited in the Protein Data Bank (PDB) with structural similarity to selected template backbones were identified using TM-align(*28*). Amino acid sequences of identified protein scaffolds were used as seeds to generate multiple sequence alignments (MSAs) using an HHBlits(*49*) search of the UniRef30 database(*50*). Resulting MSAs were used for HMMer(*51*) searches of the JGI metagenome protein sequence databases(*52*) and the Uniref100 database(*50*). HMMer search results were clustered to < 70% sequence identity using MMSeqs2(*53*) and MSAs were generated from each clustered sequence using HHBlits. AlphaFold2(*27*) was used to predict structures for each sequence using the generated MSAs. Resulting scaffolds were filtered for high confidence AlphaFold2 pLDDT scores, TMscore to the input backbone templates, and Rosetta score. Scaffolds of specific topologies were supplemented with additional AlphaFold2-predicted structures of transcription factor sequences identified from bacterial metagenomes using DeepTF(*54*). PSSMs were generated for each scaffold using PSI-Blast(*55*) and custom code for use as constraints of Rosetta design. All final scaffolds are available for download.

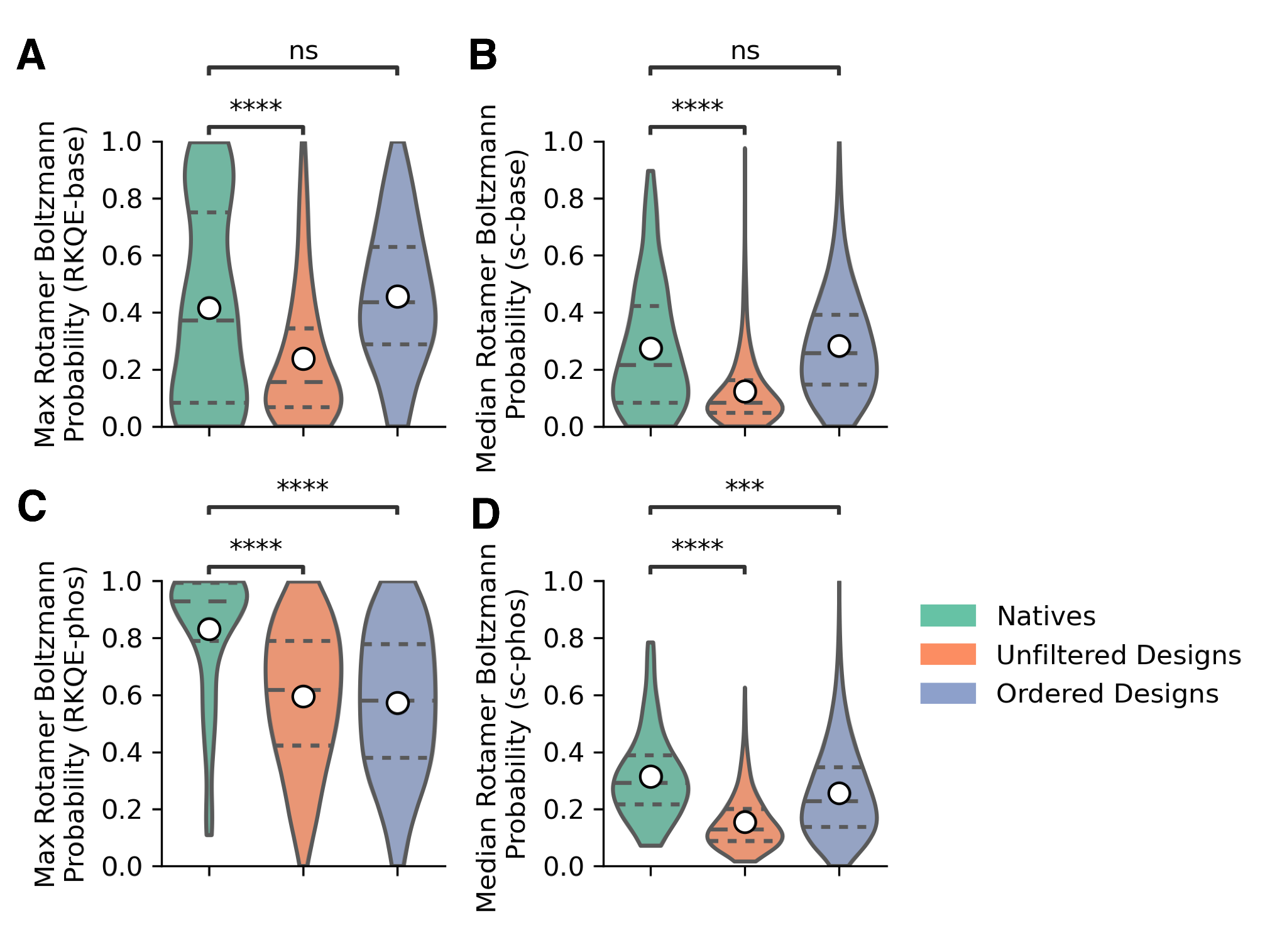

**Fig. S2. Native protein-DNA structures are enriched with highly preorganized residues forming hydrogen bonds with base atoms and the DNA phosphate backbone.** The Rosetta RotamerBoltzmannWeight metric was calculated on natives, unfiltered designs, and ordered designs as a proxy for sidechain preorganization. **Top: A,** Maximum RotamerBoltzmann probability among longer RKQE sidechains. **B,** Median RotamerBoltzmann probability for each design among all sidechains forming hydrogen bonds with bases. Among sidechains forming hydrogen bonds with base atoms, natives were significantly more preorganized than unfiltered designs. Ordered designs were filtered on the RotamerBoltzmann metric to achieve a distribution similar to natives. **Bottom: C,** Maximum RotamerBoltzmann probability among RKQE sidechains. **D,** Median RotamerBoltzmann probability among all sidechains forming hydrogen bonds with the phosphate backbone. Among sidechains forming hydrogen bonds with the phosphate backbone, both unfiltered and ordered designs were significantly less preorganized than natives. “Unfiltered Designs” were filtered by all other metrics except “Max RotamerBoltzmann Probability (RKQE-base)” and “Median RotamerBoltzmann Probability (sc-base)”. “Ordered Designs” were filtered on “Max RotamerBoltzmann Probability (RKQE-base)” or “Median RotamerBoltzmann Probability (sc-base)” and comprise all designs that were tested by yeast display in this study. Asterisks indicate p-value from a Welch’s T-test between the indicated distributions (ns: p <= 1, *: 1x10^-2^ < p <= 5x10^-2^, **: 1x10^-3^ < p <= 1x10^-2^, ***: 1x10^-4^ < p <= 1x10^-3^, ****: p <= 1x10^-4^).

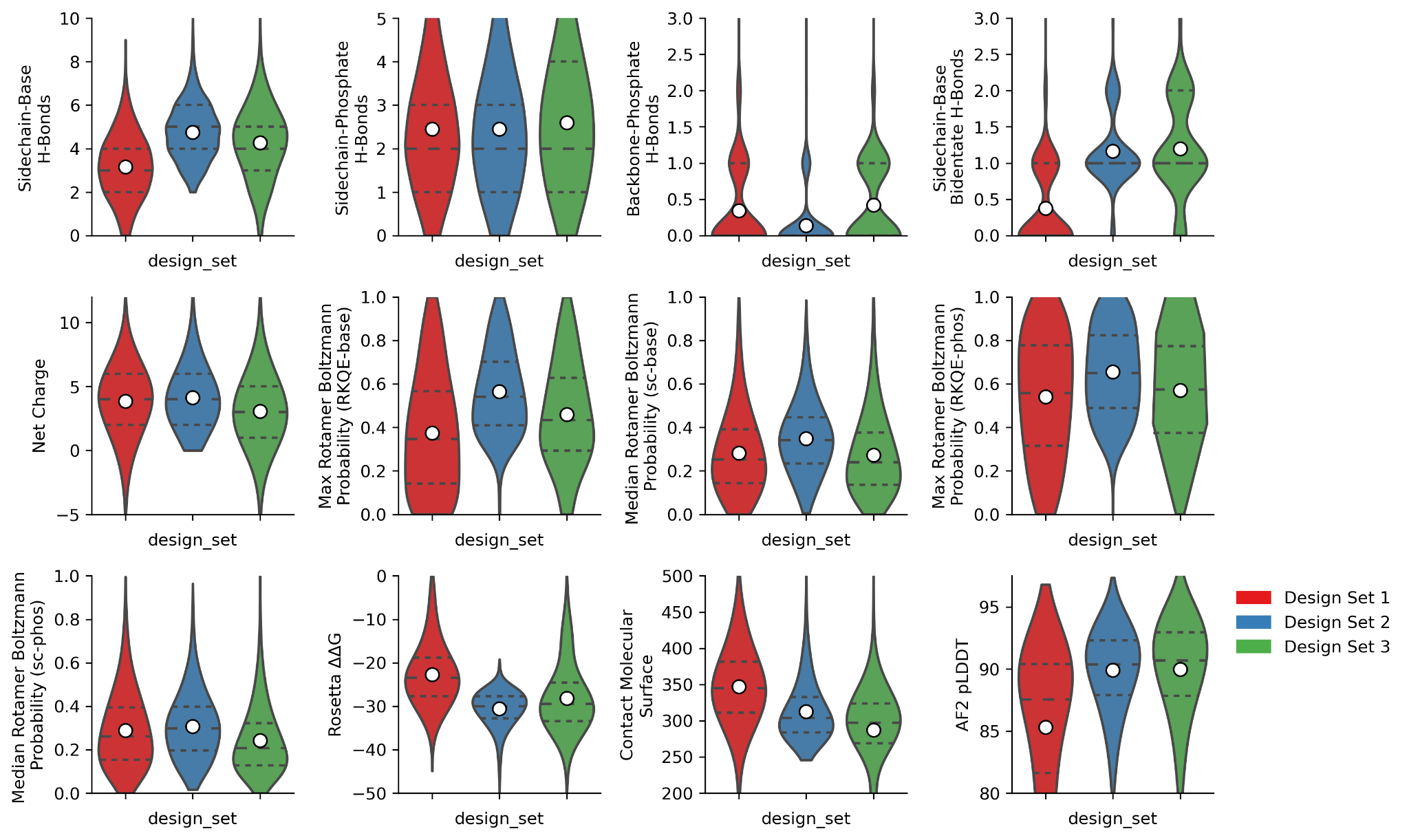
**Fig. S3. Distribution of metrics calculated on each ordered design set.** Metrics were calculated on all designs after superposition of the AlphaFold2 predicted monomer onto the initial design complexes. Design set 1 was produced using Rosetta sequence design and motif grafting, design set 2 was produced using LigandMPNN sequence design and motif grafting, and design set 3 was produced using LigandMPNN and Inpainting.

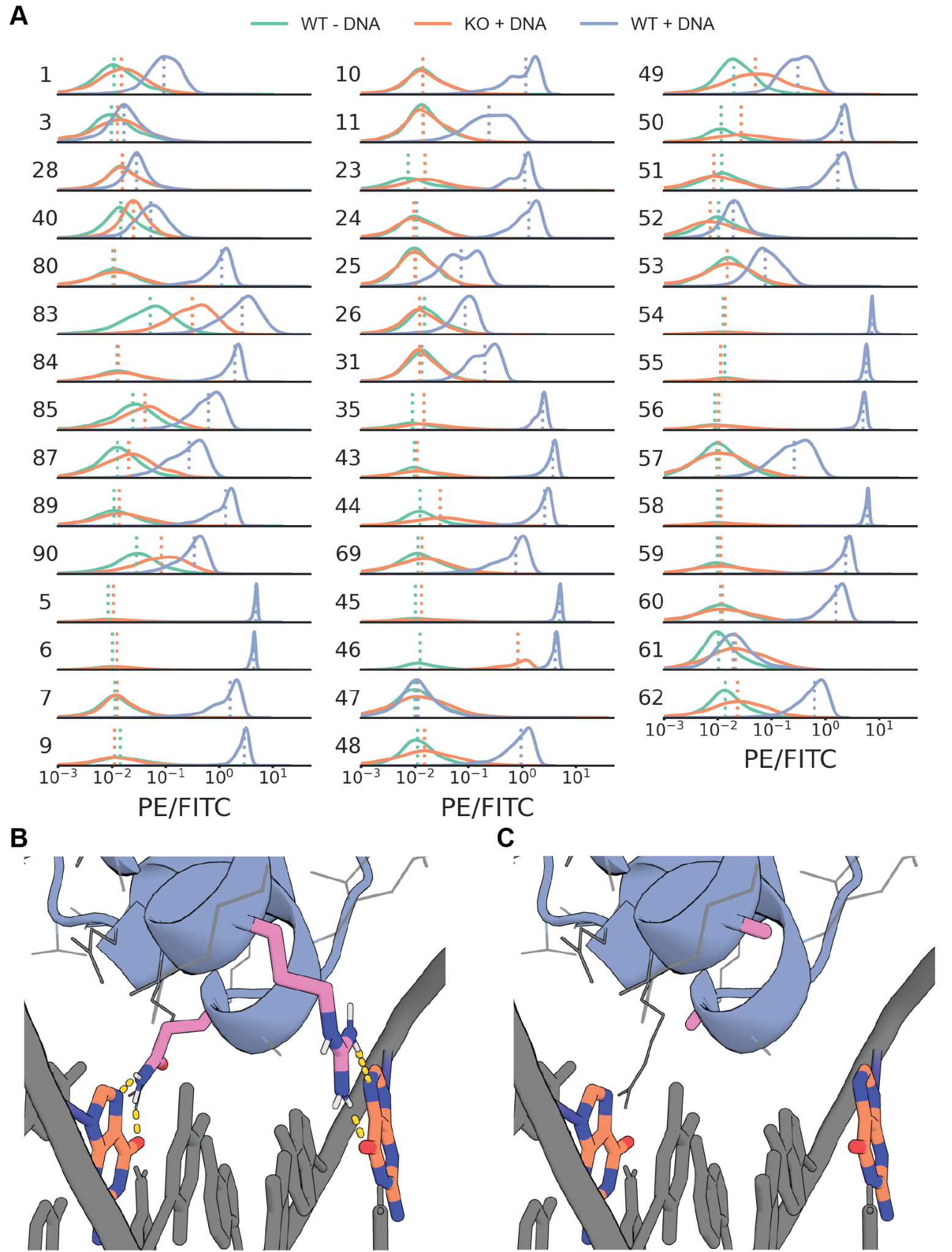

**Fig. S4. Clonal analysis of binder designs by yeast surface display confirms dsDNA-binding function. A,** Histograms of binding activity (PE/FITC) are shown for each design. Knockout sequences were created by mutating 1–3 key interface residues for base-specific contacts present in the wildtype (WT) design model (**table S1**). Samples of the WT design (WT+DNA, blue), and the knockout sequence (KO+DNA, orange) with target were analyzed after labeling with each respective dsDNA oligo at 1 µM with avidity (DBPs 7, 10, 11, 24, 25, 26, 28, 31, and 40 collected without avidity). The background signal of the wildtype design without dsDNA labeling (WT-DNA) is shown in green. Interface knockouts substantially disrupted dsDNA-binding in nearly all cases. **B,** Example (DBP43) of interface knockout of the original design model with base-specific hydrogen bonding ARG and GLN residues (pink). **C,** Model showing the two ALA substitutions (pink) of those residues.

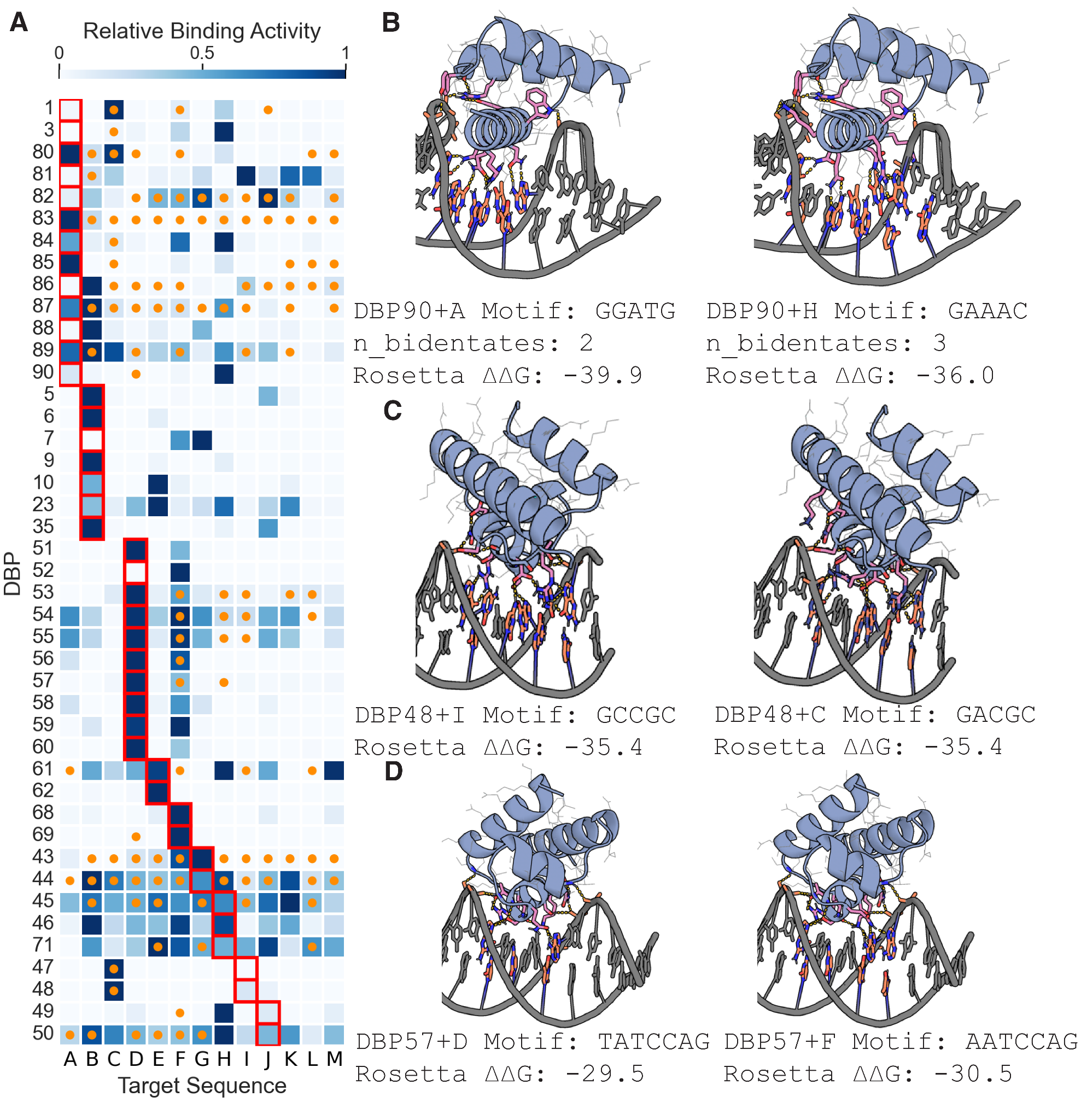

**Fig. S5. All-by-all analysis of selected designs by yeast surface display reveals preferential target binding of designs. A,** Yeast surface display relative binding activity (Normalized PE/FITC) of each design labeled at 1µM dsDNA with avidity, normalized by design row. Red squares indicate the intended target sequence for each design. Orange dots indicate target sequences containing Rosetta-predicted binding motifs. Sequences were considered potential binding targets if they had Rosetta ΔΔG less than or equal to the designed complex. DBPs 83, 85, 65, 6, 9, 35, 69, 47, 48, 51, 56, 57, 60, and 62 were considered to preferentially bind less than 3 of the 13 tested DNA target sequences, including their designed target sequence. **B,** DBP90 bound weakly to its initial design target, but strongly to an alternate target sequence (H) with slightly higher Rosetta ΔΔG but also allowed for bidentate hydrogen bonds to 3 bases. Left: DBP90+A (on-target model), Right DBP90+H (alternate target model). **C,** DBP48 bound weakly to its initially designed target sequence, but strongly to Rosetta-predicted alternative target site (D) that differed by only 1 base pair across the interface and had equivalent Rosetta ΔΔG. Left: DBP48+I (on-target model), Right DBP48+D (alternate target model). **D**, DBP57 bound strongly to its initial design target as well as an alternate target that contained an identical 6 bp stretch (ATCCAG) at the binding interface. Left: DBP57+E (on-target model), Right DBP57+C (alternate target model).

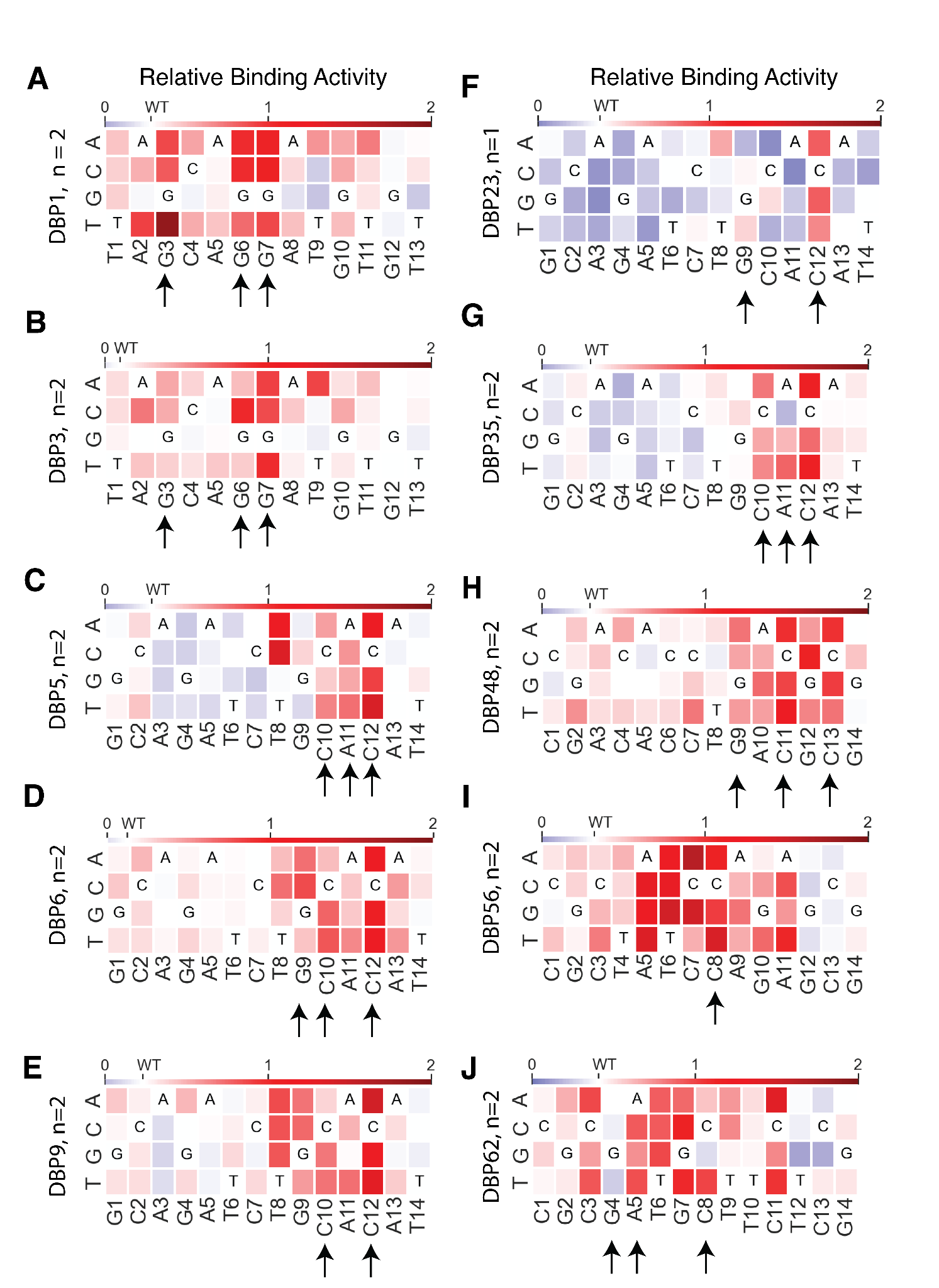

**Fig. S6. Full competition assays for all DBPs designs. A-J,** Relative binding activity (Median PE/FITC normalized to the no-competition sample) from flow cytometry analysis in yeast display competition assays for designs DBP1, DBP3, DBP5, DBP6, DBP9, DBP23, DBP35, DBP48, DBP56, and DBP62, respectively, with all possible DNA base mutations at each position of the competitor oligo. Heat maps show the mean of both replicates. Blue indicates competitor mutations where competition was stronger than with the wild-type competitor, while red indicates competitor mutations where competition was weaker. Competitor mutations where competition was weak (red) suggest incompatibility with binding to the competitor oligo. Arrows indicate base positions contacted with hydrogen bonds or hydrophobic contacts to base atoms in the design model. DBP48 was analyzed with sequence C due to its improved binding signal and nearly identical modeled binding sites. All other designs were analyzed with their designed target sequence. In several cases we observed extra specificity beyond the positions directly involved in hydrogen bonding and hydrophobic contacts. For example, DBPs 6 and 9 exhibit specificity for a 6 nucleotide stretch (TGCACA) with peripheral dependence on T8 and A13. This specificity is most likely explained by effects of shape readout that are not considered by Rosetta modeling of the designs. DBP62 appears dependent on bases peripheral to the binding site (e.g. C11).

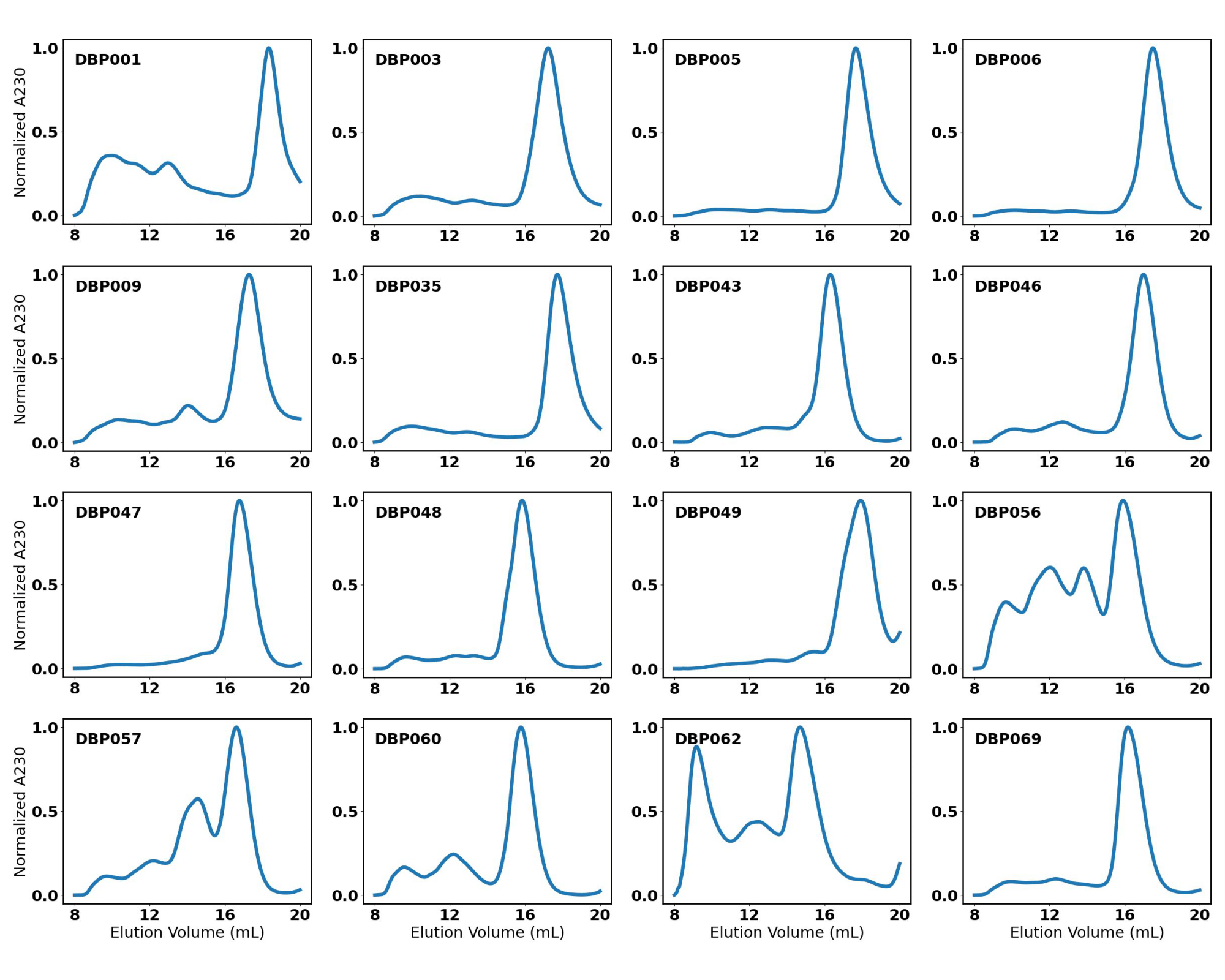

**Fig. S7. SEC traces of purified proteins.** Normalized absorbance at 230 nm of elution over a Superdex™ 75 Increase 10/300 GL column. Each plot shows a separate protein sample, following IMAC purification, with the HIS-tag attached. In every case, the highest peak, corresponding to the protein of interest, was collected and used for *in vitro* experiments.

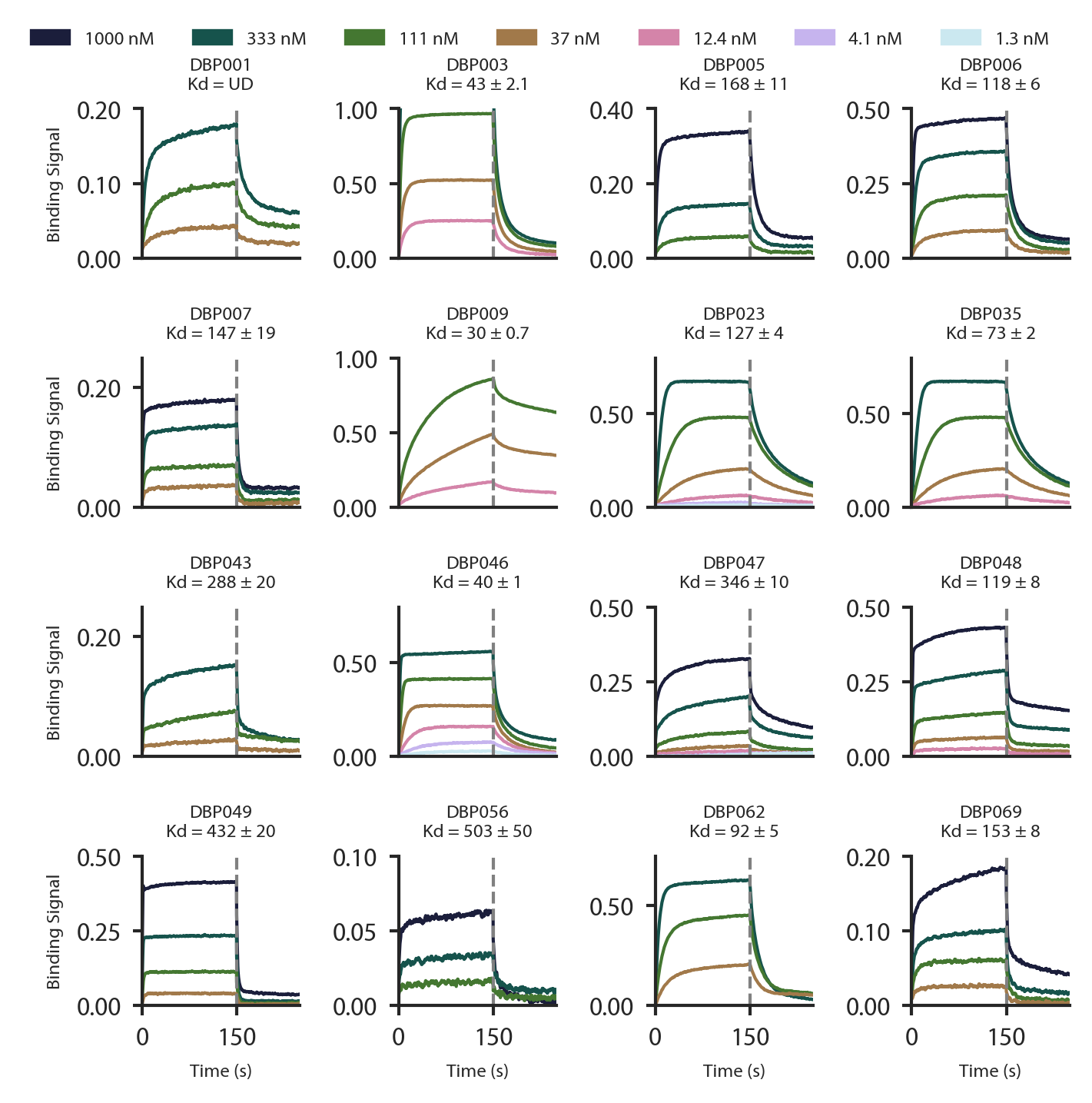

**Fig. S8. Purified designs bind their respective dsDNA targets *in vitro* by biolayer interferometry.** Binding of purified miniprotein designs to the DNA target with BLI. Each line represents biotinylated dsDNA target dilutions by ⅓. K_D_ values are indicated above each plot. UD indicates unable to determine. DBP48 was analyzed with the sequence C dsDNA target.

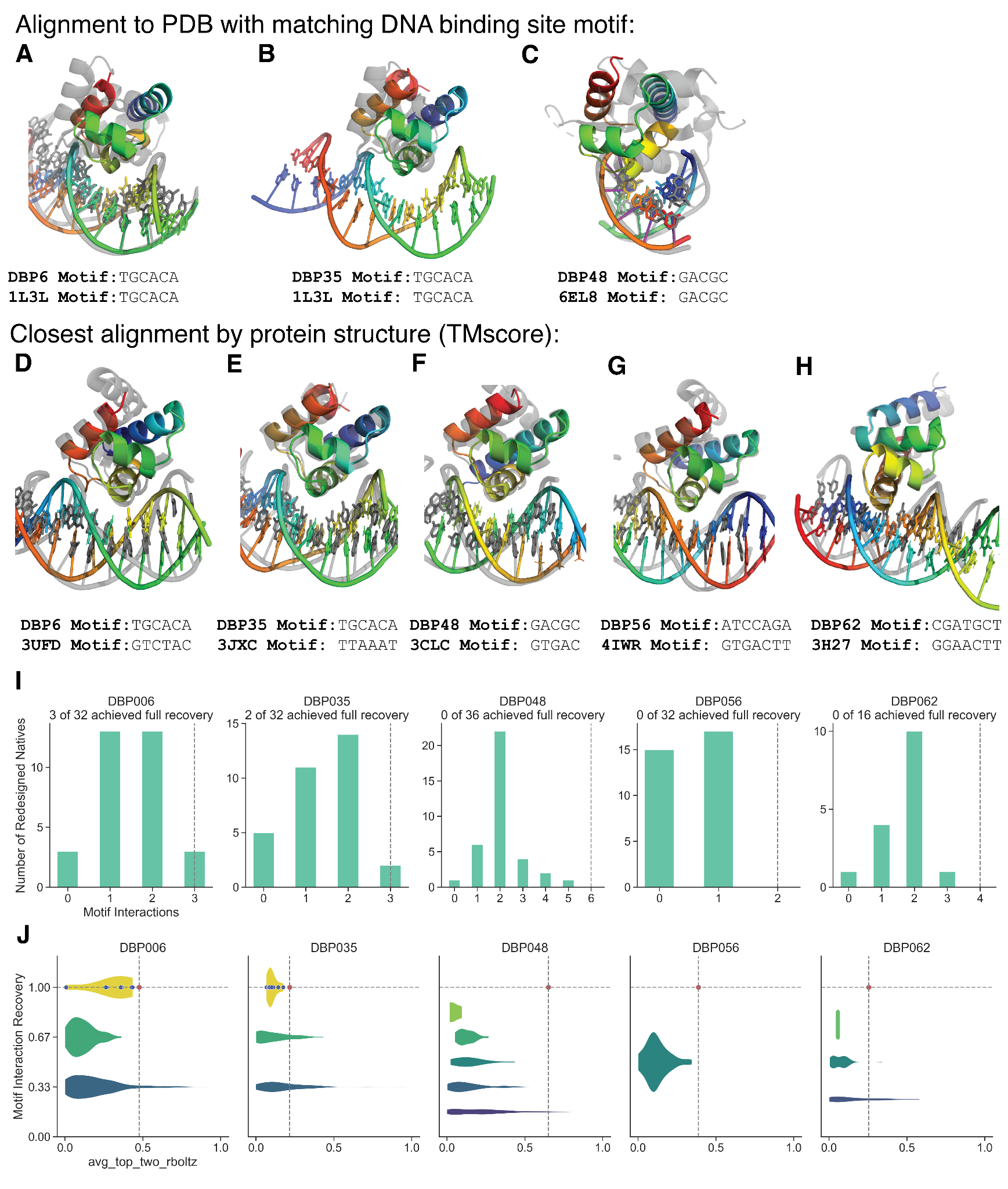

**Fig. S9. Comparison of designed DBPs with nearest native structures by target motif or protein structure. A-C,** Alignment of DBP designs to PDB structures containing an identical DNA binding site motif. DBP designs found substantially unique solutions for binding the same DNA sequence found in the native structures. Native structures are shown in gray aligned to the DBP design (colored). DNA sequence matches were found by creating a set of all contiguous DNA binding site motifs in the PDB where any atom of a protein residue was within 5 Å of an atom in the contiguous DNA sequence motif. DBP35 was designed against the DNA sequence from the crystal structure of the 1L3L PDB. **D-H**, Structural alignment of DBP designs to nearest PDB structures by TM-align. TM-align searches were performed on protein-DNA co-complex structures in the PDB to identify the nearest native protein scaffold. Nearest structures are shown in gray aligned to the DBP design (colored). Consistent with the use with native metagenome structures, most DBP designs had close matches to scaffolds in the PDB but altered docking configurations relative to DNA. The fine sampling of docking modes allows the method to identify binders to substantially unique DNA sequences. **I**, Computed statistics on native DBPs in the PDB with a TM-score to each design above 0.65 (**table S3**) redesigned in the presence of the designed DBP’s DNA target motif. Original DNA motifs in the native structures were replaced with the recognized motifs from each designed DBP. In cases where the register of the DNA in the crystal structure complex did not match the design model, we systematically slid the design motif sequence, exploring all possible offsets and generating rethreaded structures for each sequence alignment. Next, we used LigandMPNN to redesign the entire sequence of each native complex followed by sidechain relaxation using Rosetta FastRelax. To assess the resemblance between redesigned natives and designed DBP motifs, we examined whether the same amino acids formed hydrogen bonds with the same DNA base atoms (motif interaction recovery). The native redesign method was able to achieve full motif interaction recovery for DBPs 6 and 35, but not the remainder of analyzed designs. **J,** Analysis of sidechain preorganization for recovered motifs residues by average top two RotamerBoltzmann score. Violin plots show the distribution of *avg_top_two_rboltz* among recovered interacting residues for each design. Individual data points are shown for designs with full motif atom recovery (original design in red, best native redesigns in blue).

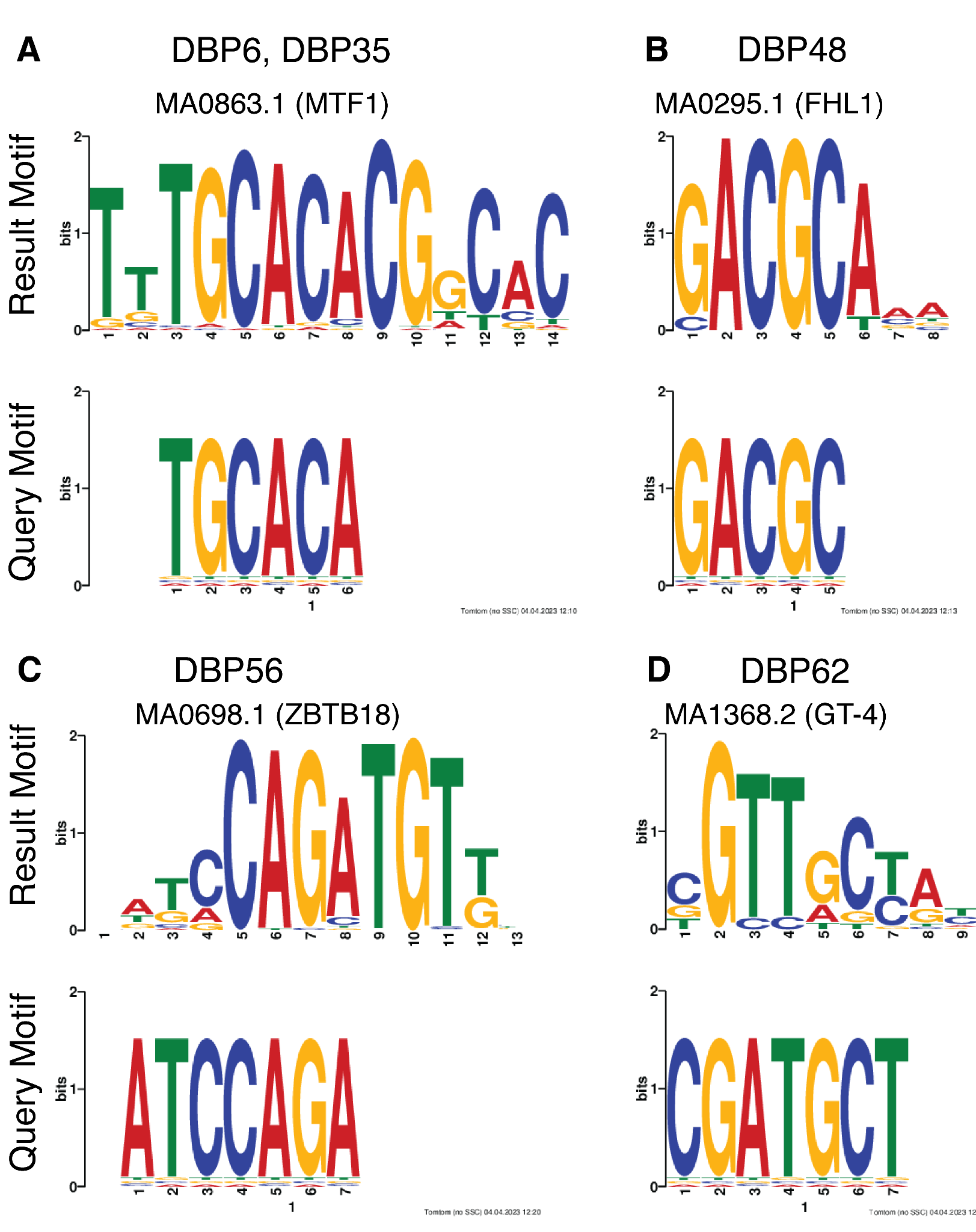

**Fig. S10. Comparison of designed DBP target motifs to JASPAR database. A-D,** Comparison of the DNA binding site motifs found through motif searches of the JASPAR non-redundant transcription binding profile database. DBPs 6, 35, and 48 were found to bind highly similar sequences as the identified search hits in the JASPAR database; however, DBPs 56 and 62 were found to bind unique sequences compared to transcription factors with known specificity profiles. DBPs 6 and 35 were designed against the DNA sequence in the PDB structure 1L3L and thus were not expected to specifically bind a novel sequence. DBP48 was designed against a novel 14 bp sequence, but the experimentally verified 5 bp binding site motif was similar to the observed specificity of the FHL1 transcription factor.

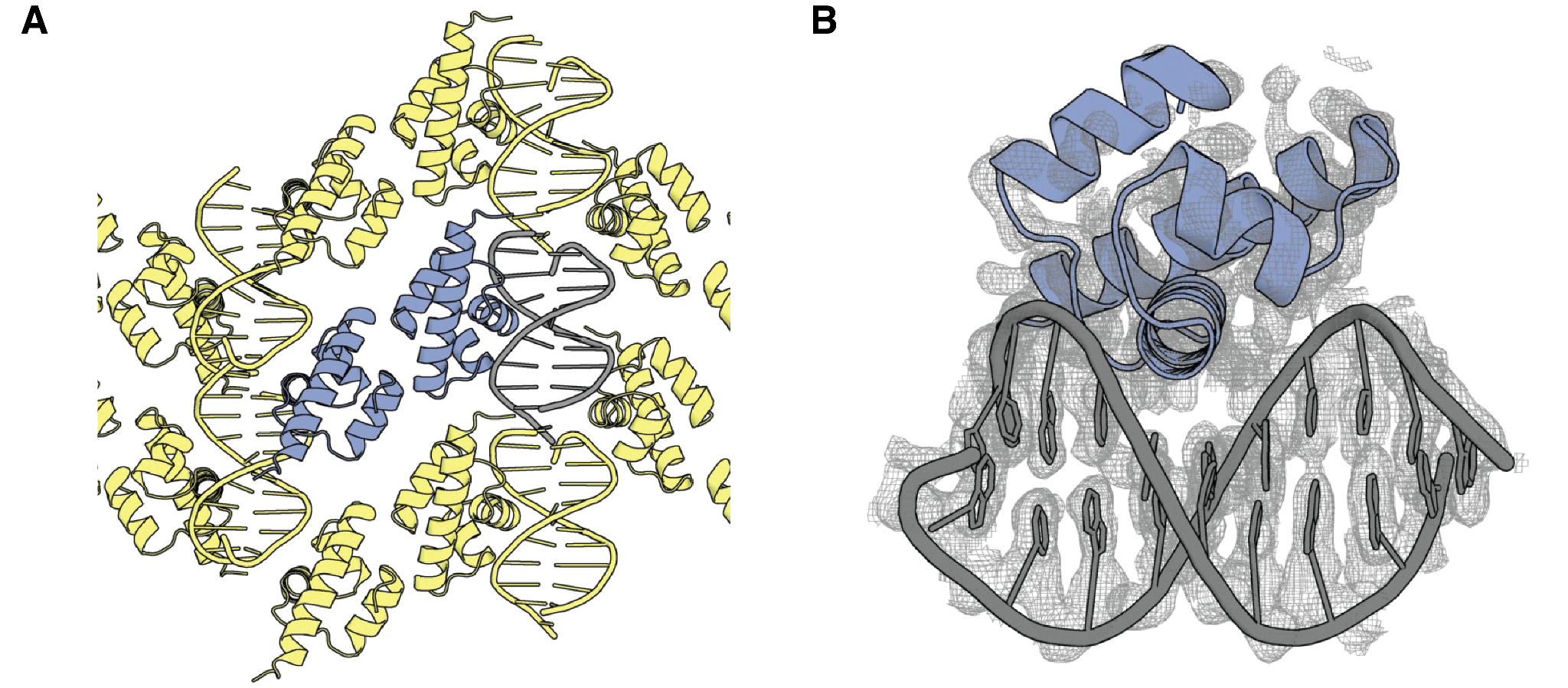

**Fig. S11. Global view of the DBP48 co-crystal structure. A,** Packing of the DBP48 co-crystal structure with asymmetric unit highlighted in blue. **B**, Global density of the DBP48 crystal structure.

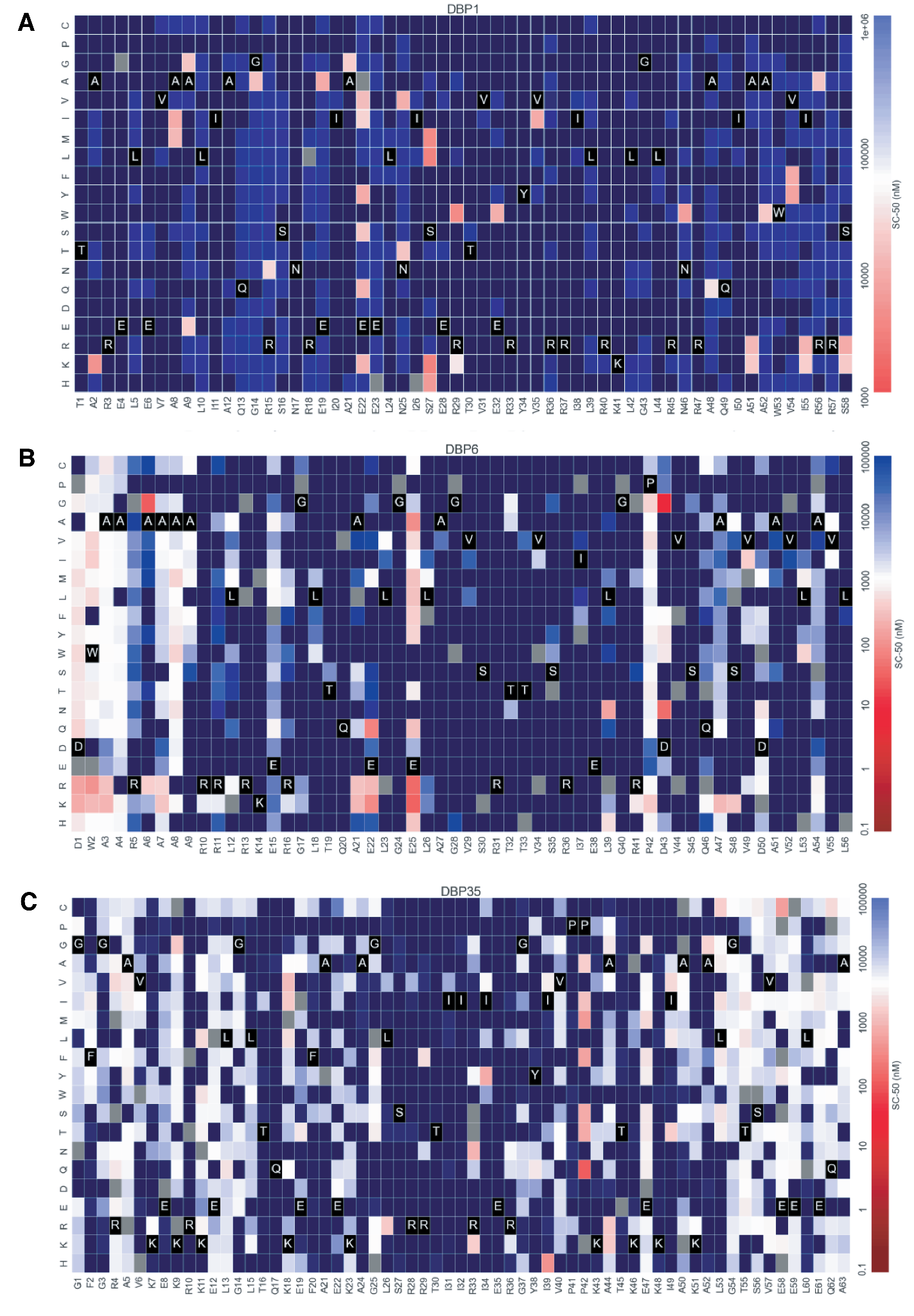

**Fig. S12. Full SSM maps for DBP1 (A), DBP6 (B), and DBP35 (C).** Heat maps representing SC-50 values for single mutations in each design. Substitutions that are heavily depleted are shown in blue, and beneficial mutations are shown in red. The depletion of most substitutions in both the binding site and the core suggest that the design models are largely correct, whereas the enriched substitutions suggest routes to improving affinity.

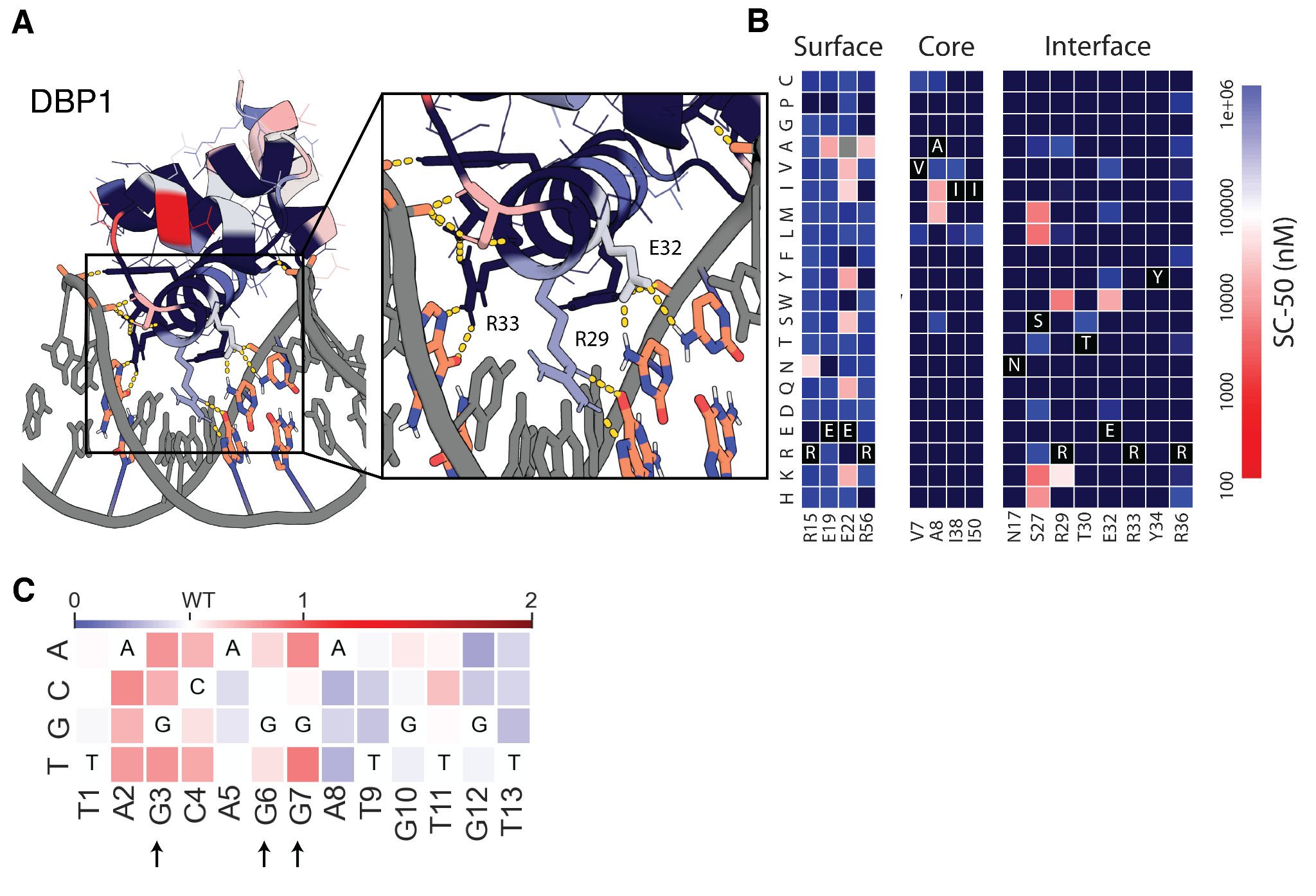

**Fig. S13. Close-up view of DBP1 SSM. A,** Design model and close-up view of DBP1-DNA interface. **B,** Heat maps representing SC-50 values for single mutations in the design surface (left), core (middle) and the designed interface (right). Substitutions that are heavily depleted are shown in blue, and beneficial mutations are shown in red. The depletion of most substitutions in both the binding site and the core suggest that the design models are largely correct, whereas the enriched substitutions suggest routes to improving affinity. For DBP1, we noticed cases where interface mutations appeared to improve binding unexpectedly. R29W and E32W mutations in particular were found to improve binding. **C,** Clonal analysis by yeast display revealed that R29W substantially disrupted specificity when compared to the original design (**fig. S6**).

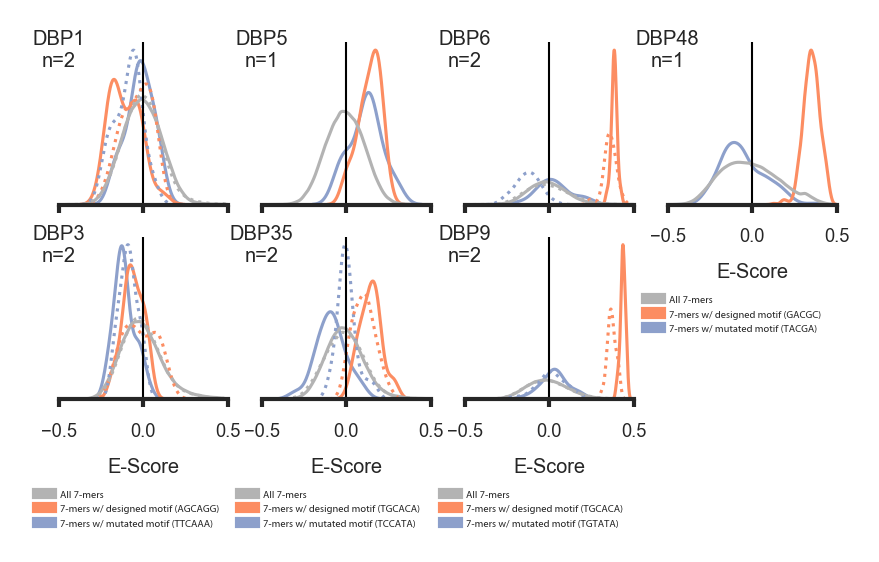

**Fig. S14. Analysis of DBPs 1, 3, 5, 6, 9, 48, and 35 with universal protein binding microarray experiments containing all 7-mers.** Solid lines represent replicate 1 while dashed lines represent replicate 2, where applicable. DBPs 6, 9, and 48 were highly specific to the intended target and the mean percentile rank of 7-mers containing the designed binding site 5-mer or 6-mer was 99.54%, 99.89%, and 97.59%, respectively. DBPs 5 and 35 were less specific to their target site, but still preferred the target motif over sequences with a mutated binding motif (designed motif percentile 86.54% and 81.88%, respectively). DBPs 1 and 3 did not appear to have a preference to the designed target site (33.19% and 46.58%, respectively).

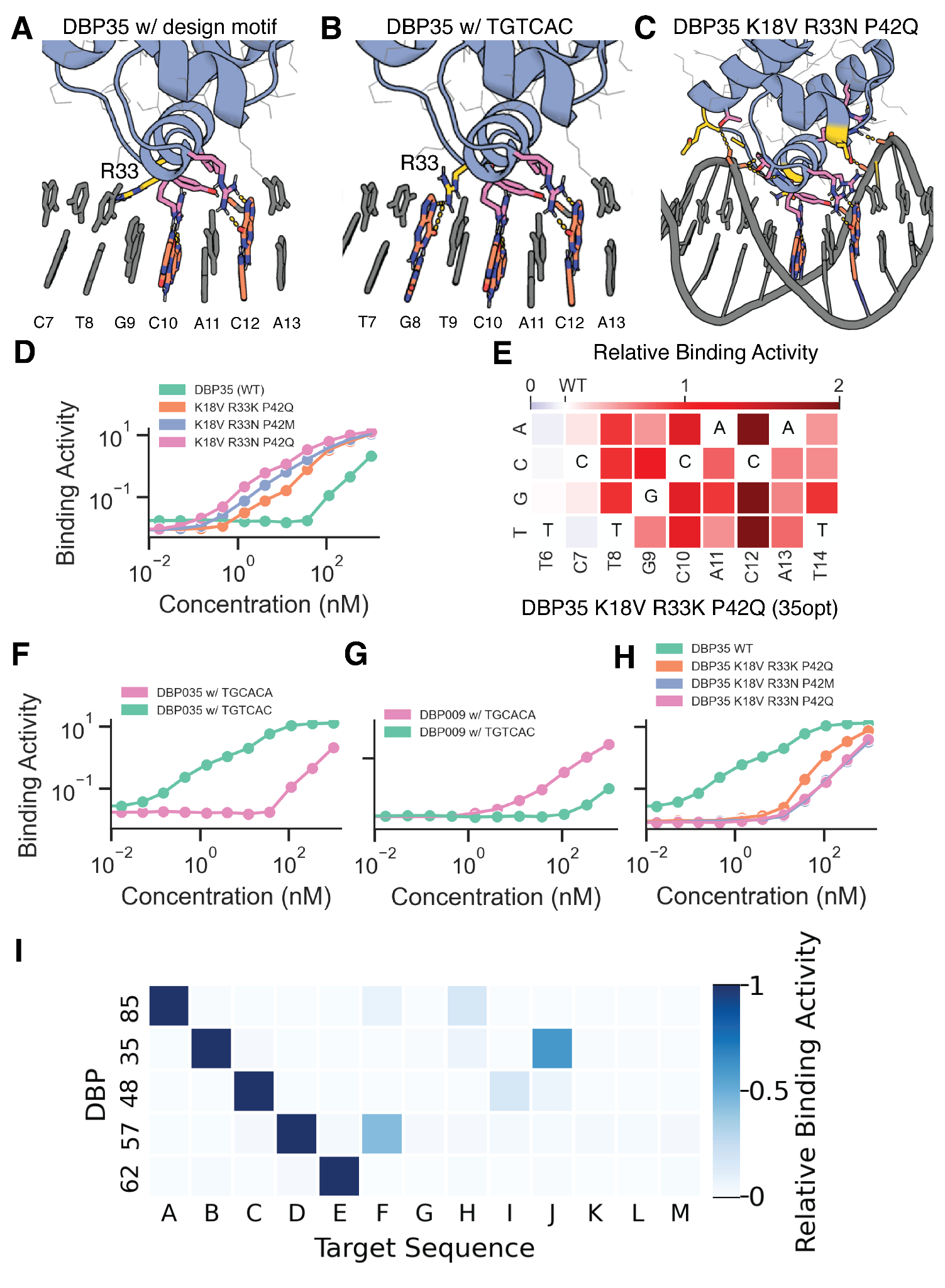

**Fig. S15. Optimizing DBP35 to disrupt off-target DNA binding.** 7-mers containing TGTCAC were enriched in designs targeting the sequence B (1L3L) dsDNA oligo in uPBM experiments. **A,** Design structure of DBP35 with R33 highlighted in yellow. **B,** Structure of DBP35 modeled with 7-mer TGTCACA shows R33 forming a potential hydrogen bond with G8. **C,** Structure of DBP35 modeled with affinity enhancing mutations K18V, R33N, and P42Q informed by SSM experiments. **D**, Binding activity (PE/FITC) from yeast display titration (without avidity) of biotinylated dsDNA target shows several orders of magnitude improvement in binding activity in DBP35 combo mutants, with binding signal detectable with dsDNA labeling below 1 nM. **E**, Relative binding activity (Normalized PE/FITC) from a yeast display competition assay of DBP35 K18V R33N P42Q showing substantial improvement in specificity over DBP35 (**Fig. 2C**). Competition assay was performed with biotinylated dsDNA target at 20 nM and competitor dsDNA at 160 nM. **F,** Yeast display titration (without avidity) showing binding activity (Median PE/FITC) of DBP35 with biotinylated dsDNA target containing designed target motif (CTGCACA) or substituted with alternative target motif (TGTCACA) shows increase in binding strength for TGTCACA over CTGCACA. **G,** Yeast display titration (without avidity) of wildtype DBP9 shows that the designed target motif is strongly preferred over the off-target sequence. **H,** Combo mutants of DBP35 show significant disruption of binding to dsDNA oligos containing the alternate TGTCAC motif by yeast display. **i**, Orthogonality matrix for 5 designed DNA binders screened by yeast display against all target sequences for which designs were made, normalized by row, at a DNA concentration of 1uM (with avidity).

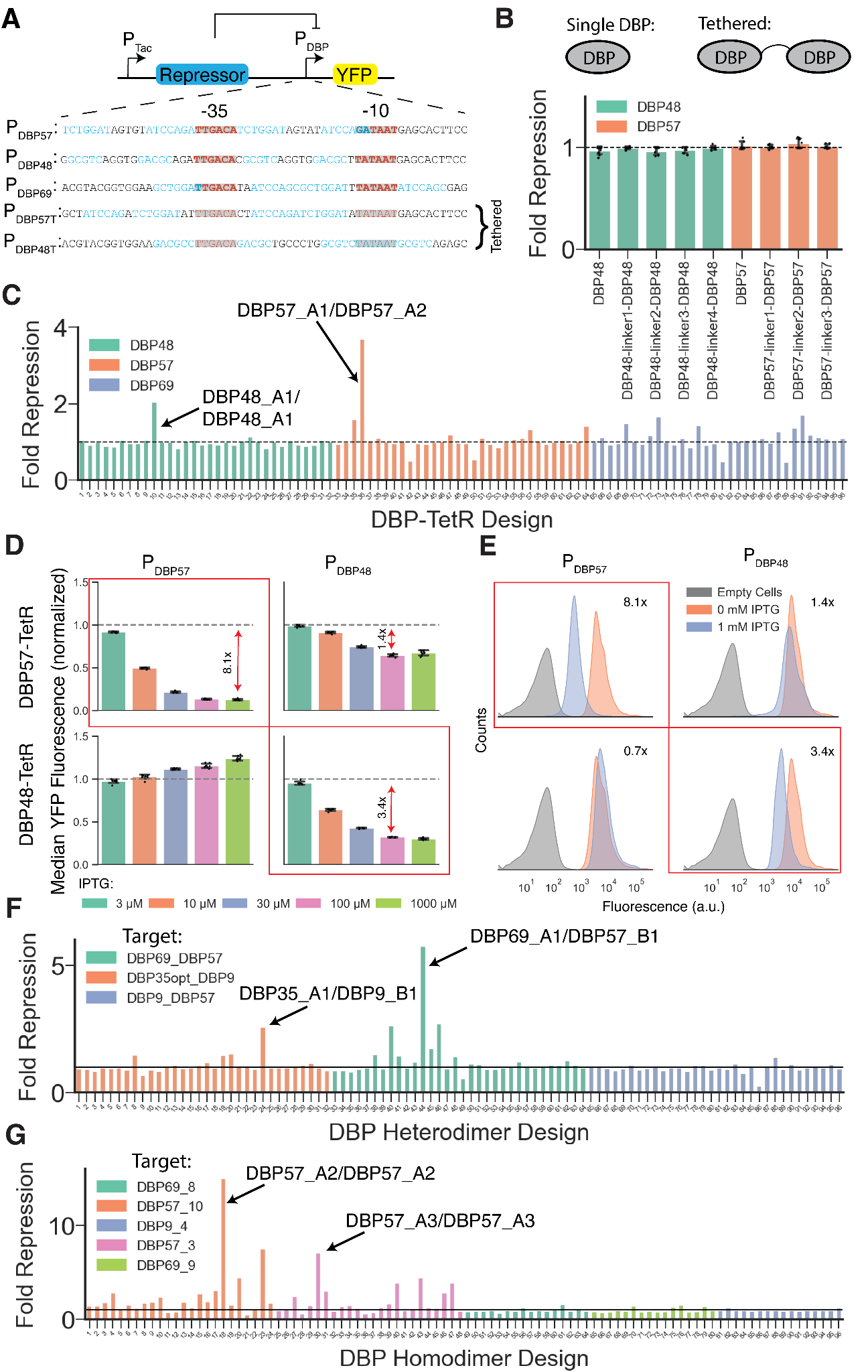

**Fig. S16. Use of DBPs to direct transcriptional repression in *E. coli.*** **A,** Vectors encoding the repressor variants were constructed with a repressor under control of the IPTG-inducible P_Tac_ promoter. A synthetic promoter containing the designed DBP binding sites (blue text) around the -10 and -35 elements (red text) was used to control expression of YFP. Alternative synthetic promoters were used for the flexibly linked DBPs (tethered) to optimize the orientation and spacing of the binding sites. **B,** Fold repression was not observed at 1 mM IPTG induction as determined by flow cytometry analysis of cells containing single DBP domains (DBP57, DBP48) and tandem linked DBP domains used as repressors. n=4. **C,** Repression screen of 96 DBP-TetR designs revealed substantial repression for at least two variants incorporating DBP57 and DBP48. Fold repression was determined for cells induced at 0.1 mM IPTG. n=1. **D,** Normalized median YFP Fluorescence from flow cytometry analysis of cells containing the successful DBP57-TetR (upper) and DBP48-TetR (lower) NOT gate circuits. **E,** Representative histograms of YFP fluorescence from *E. coli* cells transformed with DBP-TetR NOT circuits. Fold repression of YFP was ~8.1x and ~3.4x for DBP57-TetR (upper left) and DBP48-TetR (lower right) repressor variants, respectively, when encoded with their cognate promoters upon induction with 1 mM IPTG. Fold repression (mean of 8 biological replicates) is indicated in each subplot showing flow cytometry histograms of measured YFP fluorescence for empty cells (gray), uninduced cells with repressor vector (orange), and induced cells with repressor vector (1 mM IPTG, blue). **F,** Repression screen of 96 DBP heterodimer designs revealed substantial repression for 4 variants. Fold repression was determined for cells induced at 1 mM IPTG. n=1. **G,** Repression screen of 96 DBP homodimer designs revealed substantial repression for 7 variants. Fold repression was determined for cells induced at 1 mM IPTG. n=1.

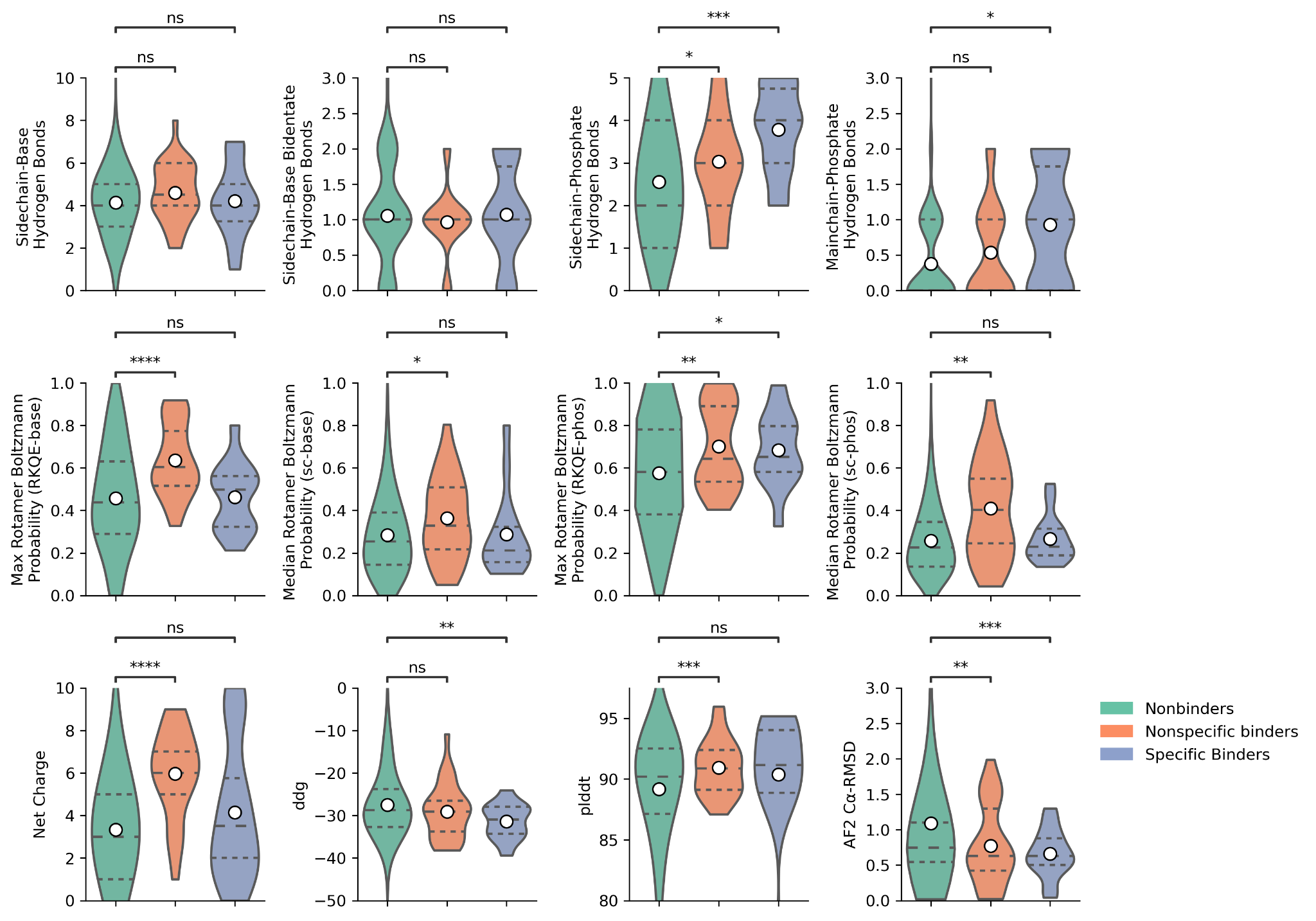

**Fig. S17. Power of computational metrics to predict binders.** Metrics were calculated on all designs broken down into nonbinders, nonspecific binders, and specific binders. Metrics associated with specific binders include number of sidechain- and backbone-phosphate hydrogen bonds, max RotamerBoltzmann probability of RKQE residues forming hydrogen bonds with the phosphate backbone, Rosetta ΔΔG, and Cα-RMSD of AlphaFold2 model to the initial design model. The different variations of the design pipeline were similarly successful in generating designs that bound their intended dsDNA oligo. While we found that generating large numbers of designs was most efficient and streamlined when using MPNN-based sequence design in conjunction with inpainting-based resampling, this approach had a lower overall success rate than the other design strategy variations, likely due to the comparatively more extensive search for suitable backbones when motif grafting with the entire scaffold library. The MPNN-based sequence design approach appeared to produce a slightly higher rate of successful designs than Rosetta-based sequence design and was also much more CPU efficient. Of the limited set of 7 designs assessed by microarray, the 4 MPNN-based designs were significantly more specific than the 2 Rosetta-based designs in the microarray context (**fig. S14**). Asterisks indicate p-value from a Welch’s T-test between the indicated distributions (ns: p <= 1, *: 1x10^-2^ < p <= 5x10^-2^, **: 1x10^-3^ < p <= 1x10^-2^, ***: 1x10^-4^ < p <= 1x10^-3^, ****: p <= 1x10^-4^).

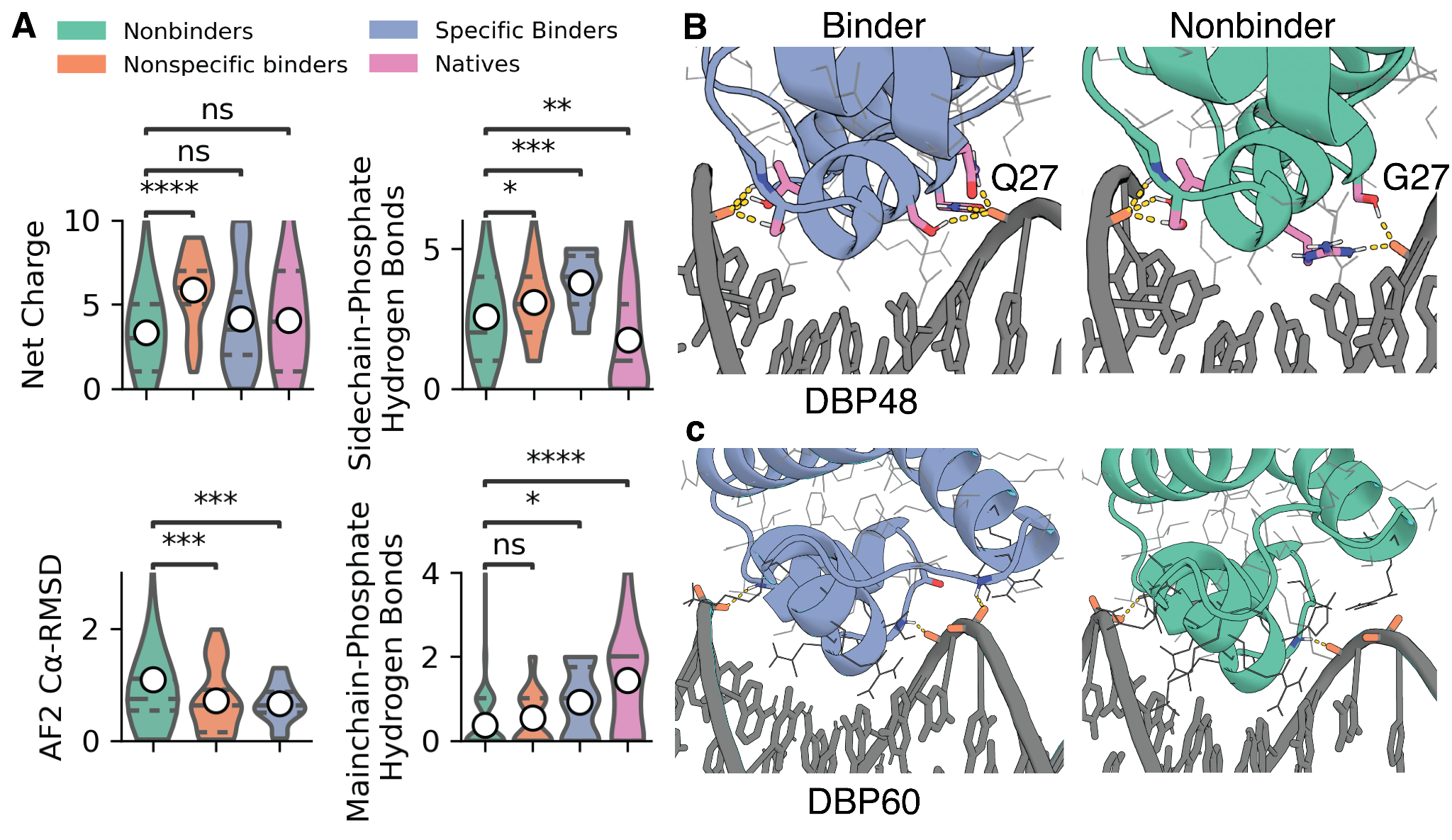

**Fig. S18. Mainchain-phosphate hydrogen bonds fix specificity and limit alternative target sites of DBP scaffolds. a,** Comparison of the statistics calculated on non-binders, nonspecific binders, specific binders, and native structures. High net charge was correlated with nonspecific binder designs, while sidechain-phosphate and mainchain-phosphate hydrogen bonds were more correlated with specific binder designs. In both cases, designs having low Cα-RMSD of the initial design model to the AlphaFold model of the protein monomers was correlated with binding. Native structures have substantially more mainchain-phosphate hydrogen bonds than even the specific designs identified. **b,** Examples of a key sidechain-phosphate hydrogen bond in the highly-specific DBP48 design and a nearly identical nonbinding design containing a Q27G that disrupted the interaction. **c,** Example of a mainchain-phosphate hydrogen bond in highly-specific DBP60 and a nearly identical nonbinding design with the terminal helix moved away from the phosphate backbone. Asterisks indicate p-value from a Welch’s T-test between the indicated distributions (ns: p <= 1, *: 1x10^-2^ < p <= 5x10^-2^, **: 1x10^-3^ < p <= 1x10^-2^, ***: 1x10^-4^ < p <= 1x10^-3^, ****: p <= 1x10^-4^).

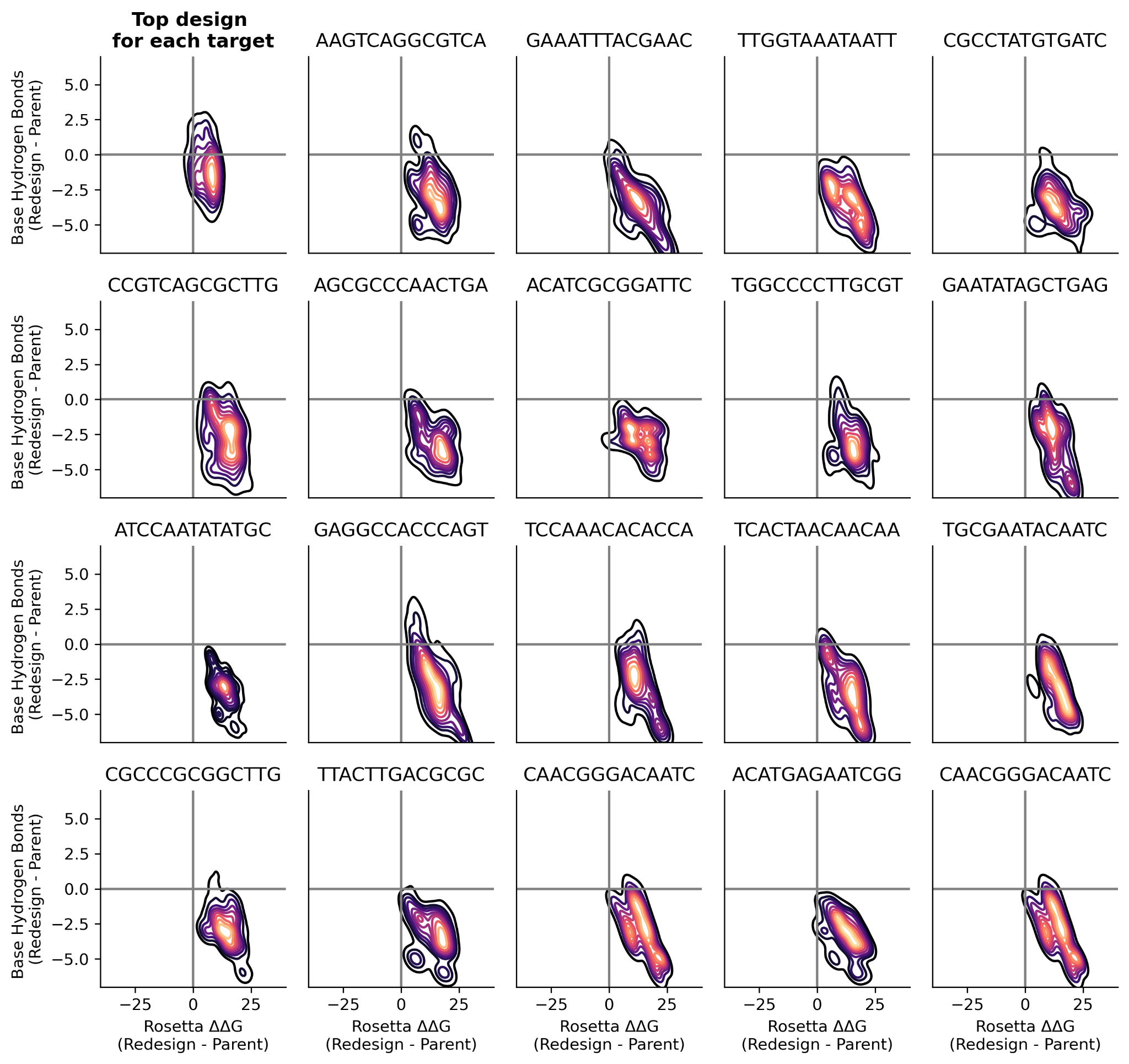

**Fig. S19. Starting dock restricts possible target sequences of DBP designs.** MPNN-redesign of successful design scaffolds against alternative target sequences demonstrates that only few alternative sequences achieve statistics comparable to the parent scaffold complex. Fourteen design scaffold complexes were redesigned against 100 randomly generated DNA sequences after rethreading onto the design complex backbone, and 20 LigandMPNN sequences were generated for each design. Top left shows a kernel-density estimate (KDE) plot of re-designs with the lowest Rosetta ΔΔG for each of the 100 target sequences. The x-axis of each plot represents the difference of Rosetta ΔΔG of each redesign from its respective parent complex, while the y-axis represents the difference in the number of base-specific hydrogen bonds. Redesigns for 2 out of 100 target sequences achieved a Rosetta ΔΔG and number of hydrogen bonds to bases comparable to their respective starting scaffold complexes. The remaining subplots show KDE plots of redesigns generated for 19 additional randomly selected sequences.

**Tables S1 to S4**

| **Design Name**  **...Closest Alignment**  **ID (% Identical)** | **Design Sequence (recognition helix)**  **...KO Mutations**  **...Aligned Sequence** |
| --- | --- |
| DBP001  HEE58516.1 (73%) | TARELEVAALIAQGRSNREIAEELNISERTVERYVRRILRKLGLRNRAQIAAWVIRRS  A AA  TAREREVAAEIAQGRSNREIAERLVLSERTVATHVANILTKLGFASRAQIAAWVV |
| DBP003  HEQ00365.1 (66%) | TKREREVLKLIAEDYGNKEIANRLNISERTVERYIRRILRKLGLKNRAELVRYAIRHG  AA AA  TEREREVLTLIAEGLSNQEIAQRLYISVKTVQTHRTHIMEKLGLHNRAELVRYAIRKG |
| DBP005  WP_028435833.1 (57%) | GFGKAVKAKRAELGLTQAEFAERAGLSRRTIIRIEQGKVKATSTTAEKIAAALGTTVQELEQA  AA  RAERGWTYDELAERSGLARRTLIEIEQGRTVGTLRTWHAIAHALGAPVGEL |
| DBP006  WP_073716028.1 (68%) | DWAARAAAARRLRKERGLTQAELGELAGVSRTTVSRIELGRPDVSQASVDAVLAVL  A A  LRKERGMTQADLGTLAGVSRQTIVSIEKGHFDPS |
| DBP007  WP_107251623.1 (64%) | PLAELGKAIREARKKKGLTQEEVAKAAGVSRATVQRLELGKAKSIAPEKLAAIAKVVGL  A AA  LGDRIRERRMRQNLTQEKLAEAAGVSRDTVQRIERG |
| DBP009  WP_095597620.1 (50%) | DWERRCAYARRARKELGLTQAELGELAGVSRTTVSRIERGKPDVSEASVEAVLAVL  AA A  ANVRRRRLELGLSQEELAEAAGVHRTYVGMLERGEKNVTIYNIERI |
| DBP010  WP_124270774.1 (64%) | PLAEIGRAIKEARKRRGLTQAEVAEAAGVSRATVQRLELGKAKSIAPEKLAAIARVVGL  T TT  LGRAVREARIRQGMTQTQLAEVAGVSRATLQNLERG |
| DBP011  WP_006546351.1 (64%) | VGEWVKRKRKEKGLTQEELAKLLGTSRATVQRIELGKKAPTPEQLERARRILEE  A AA  RKEKGLSQQELAKLVGVSRNTISSIETGQYCPT |
| DBP023  WP_005182224.1 (61%) | PLAELGKAIKEARKRKGLTQAEVAKAAGVSRATVQRLELGKAKSIRPDKLRAILEVVGL  A AA  LGTAIREARKAKGLTQAQVAEAAGLARSTVIEVERGQ-KSLSSDAL |
| DBP024  OIP90164.1 (61%) | PLAELGRAIREARRRRGLTQEEVARAAGVSRATVQRLELGKAKRIRPEKLAAIARVVGL  T TT  LFALGRSIREARKRRGLTQAEVAKAVGIGRAALSRLEGGVIREI |
| DBP025  HGZ71070.1 (56%) | PLAEIGKAIKEARKEKGLTQEEVAKAAGVSRATVQRLELGKAKSMRPEKLAAIAKVVGL  T TT  LLELGQTIKELRKERKLSQEELAKQSNISRATLSKLENG |
| DBP026  HDY16147.1 (49%) | DP----ILELLLEGEHTATELMRRLGLSYRTVRSRLRSLVRQGIIGYRHTGRVVYYVRDPERVRELMAR  A A  DPTRVRIVELLLEGEKNVSELVEALGMQQGRVSSHLACLKWCGFIGTRREGKFVYYRVTDERVRELM |
| DBP028  PYL53864.1 (45%) | SPLVLAILEGVARGRTPAEIAKELGVSRRTVQNILQYLRRKHKLSL---EELVPFARRVLAAR  A A A  LAVFSMIAAGRRPGEIAKELGISRKTVNTHFEHI--KHKMGYANAEELKHGARELLGS |
| DBP031  WP_081958439.1 (58%) | LAAEIKRLRREAGLTQRELAERMGVSRYTVQRYELGKRTPSPEELERILAALGV  A  IKELRRDNGLTQKELANAMGVTASMIGQYETGIRTPKYETLERIAKALSV |
| DBP035  WP_143902275.1 (55%) | GFGRAVKEKRKELGLTQKEFAEKAGLSRRTIIRIERGYIVPPKATKEKIAKALGTSVEELEQA  AA  REHLGLTQAEVARRAGMSQAALSQIESGEHKPRKATREKLAKALGITVEQL |
| DBP040  WP_167784858.1 (59%) | SAERHFLAVAEAGSYRRAAEILGVRRDTVRRAVLRIERKLGAPLFRREPV-LTLTPLGRELYERLQ  A A  RHFLAVARHGSTLAAAPVLGVDPSTVNRRVRELEKRLQAVLFRREPSGYTLTELGRSL |
| DBP043  HAS56005.1 (56%) | MEELKKLLESGDPKLQGEAVKKIREKLGLTQREFGKKIGVGQAKVSRIEAGKIKLTPELKEKILE  A A  YSEAIKKLRNKLILTQTEFGEVFGVSFATVNRWEAGKFKPTTKLKRKL |
| DBP044  RLI39306.1 (49%) | KEERVLAALEAHPWSTLAEIAELTGLSRSTVSRILSRLRKEGKCDSRREGRKVRYWLVRR  AA A  KILAALEERPGLTVAQLAELTGLSRSSVRLHLASLRREGVVD--RRGKRPYTWFL |
| DBP045  WP_129869568.1 (60%) | DREIEEFRREVRERMAEQGLTQADLARRSGLSRNTISRFLRGKTRPTPATVEAIRRALGLPA  AA A  LTQEQLAERSGLSRNTIIRIERGNTNPSMKTLDRIARAMG |
| DBP046  WP_137097520.1 (50%) | RRELSPLARARKAAGLTQRELAEKAGVGTATISRIERGRRPFSRLPPEKQERIAEILGVSVAELE  G G  LARFRKARDLSQEQLAAAASVGVDTVARIEQGKRSTCR--PTTLQRLADALGVSVENL |
| DBP047  WP_145853098.1 (60%) | MTPEERERAKEIGREIRELRRERGLTQRELADLLGVSRSTVSDIESGRRLPSEELLRRIREILGV  A A A  VREQRRHRGLSQAELADALGVSRQTVISIESGRYLPSLPLAFTIARFF |
| DBP048  WP_130029327.1 (62%) | MTPEEIAEAKRIGKEVKERRKELGLTQRELAEKLGVSRSTVSDIENGRRLPSEELLKKIKEILGV  A A A  GIDLKRFRKELGLKQRELAEKLAIERSLVSKIESGKRLISKELEKKIIEALHI |
| DBP049  WP_005543476.1 (39%) | MEELEERILALLREEWPRGLGAAEIARRLGVPRSKVRTALRRLVAEGRVRVVRGRYSRYVAVEP  AA A  EMEGKIIVSLSEQ--GAMTGYEVAKKLGVSRSNVYAALQRLVDQGVILHRNGDPSQYTALQP |
| DBP050  WP_056730679.1 (46%) | GNPRKEKILEALCRGPRTSTEIAREIGVSTRTAAGLLQGLVRQGLARPRRRGRRVYYELADPSIC  A G A A  GSGRRAEIIDVLAQGERAVEEISMEIGQSTANTSHHLQQLLRSGLVRTRRAGTRIYYRLAGPSVT |
| DBP051  RWE19644.1 (62%) | MEADPAVVFGRRLRAARRAKGLTQAELAERAGLGQGTISRYEKGRTLPSPEQVEKLLAAL  A A  ARDARGLTQAELADRMGVGQGTLSKYETGFSDPPREFVEDLAAEL |
| DBP052  WP_041440254.1 (65%) | RPLTPAEVFGRELRRLRRAAGLTQAELAERAGIGQGTVSRYEHGRRLPSPEEQERLLAAL  A A  LGRELRRLRAAAGLSGAELARRAGVPQPTVSRVETGRRVADPEVVVRLLGVL |
| DBP053  NLJ00870.1 (63%) | RKLSPYERFGREIKERRKEAGLTQAELAELAGVGQATVSRIEKGEKVSPEILEKIREALEKA  G G  MERFGNIIKERRKELSITQRELAALAGIGINTLTKIERGE |
| DBP054  WP_111307537.1 (62%) | RVKTPFERFGEFVKRERAKAGLTQAELAKLAGVGQSTVSRIEKGKKCSPELREKIVKALKAV  A A  LGTFIKDHRKKAGLTQLELANLAGVGKTTVFDIEKNKET |
| DBP055  WP_062332060.1 (64%) | RKKSPLEIIGERIKKERKELGLTQAELAKLAGIGQSTVSRIEKGEKCSQRIIEKIFKALAAV  A A  EILGKRLKQQRLYQNLTQVELAQRAGIGLSTVSRIESGE |
| DBP056  SHL87335.1 (37%) | PPPTPFEVAGARIKEERAKLGLTQAELAKVAGVGQATVSRIEKGRKCSWELIEKIFEALKKV  G G  PSILKEMGSRLKEYRLRKGLMQSELAESAGVGVGTIAKMERGGAVSVQILLSVLRSL |
| DBP057  WP_016182689.1 (64%) | MVLTPMERIGEFIKRARREAGLTQRELAELAGVGQSTVSRIEKGEKCSPELVEKILEALRKV  AA A  KARENAGLTQRELAELANVPQSTISRIEKGYNTSFDTLSKIASALGK |
| DBP058  WP_099045111.1 (40%) | AWTGEQLREFRKKLGLSQREFGELLGVGQSTVSRVEHG---------GELGPATRARLQARVDELVAEYKASQ  AA A  KVHDLRENMGLSQREFASLIGKPQSTIARIESGSMNASTKMLSEIAQATNQRL |
| DBP059  WP_044547039.1 (60%) | MKELGKKIKERRKKLGLTQAQLSELSGVGQGTISRLEQGRGNPSPKILEKIEKVLKELEK  A A  MKEIGKIIKERRKSLKVNQLELSELAGVGINTLVAIERGEGNP |
| DBP060  WP_155241367.1 (54%) | DIEKIAKAVKELREELGLTQAEFAKKIGIGQGTLSRFEKGGVLSPKTM-ERLLKALEKEFGFDVKK  A A  AVLQLREELGLTQAELADRAGLKQPAVARFEAGGTMPTIPMLERFAEALE |
| DBP061  No hits found | GAKEKLWEFLLELAKKGLPFKLPSAEEIARRLGVRRRTVIGQLQSFVREGRIKLKRGVVYSVNE  A A A |
| DBP062  MBE06805.1 (57%) | KEELEKLLKIIESLPKKFREVIILKFVEGLSYTEIAERLGVSRGAVYSRLRSALKKIEEALKK  A A A  DKMTIAINSLSEDHREVILLKEVEGLSYQEIAESMGSSIGTVMSRLYYARKKLQETLK |
| DBP068  NDC22157.1 (50%) | PLSGKELGELIKKYRDEKGLTQAEFAKLAGLGQGTISRLEKGVDRNGKEYHPGEEIREKVLKAIA  n/a  SNQKIGQLISQIRQERGLTQSEFAKLLGTSQSAVNRMEHG |
| DBP069  HAG83907.1 (69%) | MKEEGRKLKELRERLGLTQAELAEALGLGQSTISRLERGRKEISPEVWEKALALLE  n/a  LKELREYLGLTQEELGQALGITASTISRYERGQHQ |
| DBP071  PWM01863.1 (55%) | NTELLKQKIKEKGLSREEVAKKLGISRNTLTQKILGHRKFSPEQIEILKELLGLSEEEVKEIFFP  n/a  SREEVAAALGVTANALTNYECGIRRISLEQILILAILYGISTEEVIE |
| DBP080  WP_136053633.1 (62%) | ATAAQRWRLSPRETEVLELLINGYTNKEIASALNVSVRTVEVHIRRVLRKANVRRRVELVAKYYG  A AA  LTAREVEVAMLAVAGAANRDIAAALNVSVRTVEVHLGRVFAKLDVRRRVELTA |
| DBP081  WP_020512267.1 (71%) | TPREREVLNLLAQGYSNREIAERLNISEKTVKNYVRNILRKLGVRNRVEAVRWWLAVR  n/a  TEREREVLTLLARGSSNAEIAERLVVTEATVKTHVSAILRKLGVRDRVQAV |
| DBP082  WP_094947301.1 (71%) | NRIDSLSPREREVLRLIAQGYNNKEIAEQLNISEKTVKVHVRRILRKLNVHNRAELVNLK  n/a  LTDRERDVLRLIATGLSNKQIAVQLFISEETVKVHIRNLLRKLNVHSR |
| DBP083  WP_072606916.1 (65%) | PREREILRLLAEGKNAWEIAQILNISVRTVRNHLRNAMRKLGARNRVQAVARALRLG  A A  PRETECLKWAAEGKSEWEISQILGISEHTSEKHLLNAKRKLGAVNRVHAVAEAIRLG |
| DBP084  HEQ00365.1 (66%) | TKREREVLKLIAEDYGNKEIANRLNISERTVERYIRRILRKLGLKNRAELVRYAIRHG  AA AA  TEREREVLTLIAEGLSNQEIAQRLYISVKTVQTHRTHIMEKLGLHNRAELVRYAIRKG |
| DBP085  WP_132296588.1 (69%) | TLSQLTPQEMRIARLASEGMPNREIATRLFISPRTVEWHLRRAMRKLGVRNRTQMARRIDTRL  AA  TKNQLTAQESQIARLARDGFSNPEIATRLFISPRTVEWHLRKVFTKLGIRSR |
| DBP086  WP_088187152.1 (63%) | TKREAEVLELLSRGRSNKEIASILHISVRTVEWYIRRILRKLGVKNRVEAVRTAKAQG  n/a  TKRETEIIRLVSRGLHNKEIASVLGISVRTVEFHVSNILLKLEVSTRLEAV |
| DBP087  WP_157440657.1 (71%) | GVERLTPREKRVAHLAAQGLTNREIAEALHISPRAVENHLRRILRKLGIRRRRELPEALGE  A A  LTTREREVAELAAQGLNNREIAELLHISTRTVETHIRRLLPKLGLRRRAQLP |
| DBP088  HBK85480.1 (71%) | PREMEVLNLMAQGYNNKEIAARLGISEKTVKNHVRRILRKLGVRNRVQAVIIAQRNG  n/a  PRELEVLRLMAQGYTNLRIATELFISEKTVKNHVSSILRKLGAKDRTQAVV |
| DBP089  EHN12857.1 (67%) | SPAAFDKLTARELAVARLVAQGLPNREIAAALHISPRAVEAHLRKIYRKLGIRRRRELAALLA  A A  DDLTAREREVCDLVAGGATNREVAAALFLSPRTVEHHLRGAYRKLGVRSRSELA |
| DBP090  XP_005452710.1 (62%) | NKYQLSLLESAFQSNRYPDISQRATLASQTGLPERRIKIWFQNRRQRWKRKK  AA A  SKAQLSQLESAFSATPYPDINGRKTLASLTGLPESKIQVWFQNRRARFFKTK |

**Table S1.** Sequences of DNA binder designs, corresponding interface knockout positions, and aligned best hit of blastp search of the NCBI standard non-redundant protein database. DNA binder designs are annotated with recognition helix (underlined). Sequence ID and % identical of best hits from a blastp search against the NCBI standard non-redundant protein database with E-values > 0.01 and coverage > 60% are indicated in the ‘Name’ and ‘Protein Sequence’ columns, respectively. The recognition helix of each binder design is underlined.

| **Name (PDB ID)** | **Sequence** |
| --- | --- |
| A (1YO5) | TAGCAGGATGTGT |
| B (1L3L) | GCAGATCTGCACATC |
| C | CGACACCTGACGCG |
| D | CGCTATCCAGAGCG |
| E | CGCGATGCTTCTCG |
| F | CGGCTGGATTACCG |
| G | CGAGAACATAGTCG |
| H | CGGGGAAACGCCCG |
| I | CGCCCAAAGCCGCG |
| J | CGGAGGTAATGACG |
| K | CGCACCGACTCACG |
| L | CGGCCCTTTGCGCG |
| M | CGCCGTTAGTGTCG |

**Table S2. Sequences used as dsDNA targets.** Sequences were ordered from IDT as dsDNA duplex oligos with a 5’ biotin modification (‘/5biosg/’) on one strand.

| **Designed DBP** | **Redesigned PDB IDs** |
| --- | --- |
| DBP 6 | 3UFD, 4IWR, 3CLC, 4X4I, 4IVZ, 4YG1, 1PER, 3JXC, 2R1J, 3CRO, 3L1P, 2OR1, 5JUB, 2XSD, 1O4X, 1E3O, 1GT0, 1HF0, 1CQT, 6CHV, 1OCT, 3D1N, 2H8R, 2D5V, 1AU7, 3O9X, 3ZHM, 1IC8, 2O4A, 4JCY, 4JQD, 3ZKC |
| DBP 35 | 3JXC, 2R1J, 4IWR, 3CRO, 3CLC, 5JUB, 1PER, 4X4I, 3UFD, 2OR1, 4IVZ, 4JCY, 4JQD, 3ZKC, 3ZHM, 6CHV, 1LMB, 4Z5C, 4YG1, 3BDN, 1GT0, 2XSD, 3L1P, 3D1N, 1HF0, 1CQT, 1O4X, 1E3O, 2H8R, 1OCT, 1IC8, 1AU7 |
| DBP 48 | 3CLC, 4IVZ, 3UFD, 4IWR, 4X4F, 2R1J, 1LMB, 4JCX, 5JUB, 3BDN, 1PER, 6CHV, 4JQD, 2OR1, 1RIO, 4PU4, 3L1P, 3ZKC, 3CRO, 1GT0, 1E3O, 1O4X, 4Z5C, 4YG1, 1HF0, 2XSD, 5K98, 1AU7, 3ZHM, 3O9X, 3D1N, 1IC8, 1CQT, 2H8R, 1OCT, 1KQQ |
| DBP 57 | 4IWR, 3UFD, 3CLC, 4X4G, 4Z5D, 4IVZ, 4YG1, 1GT0, 2XSD, 1HF0, 3L1P, 3D1N, 1E3O, 1O4X, 1CQT, 1OCT, 2H8R, 3CRO, 3JXD, 1IC8, 1AU7, 2R1J, 5JUB, 1PER, 2OR1, 6CHV, 2D5V, 3ZHM, 3ZKC, 4JCY, 4JQD, 2O4A |
| DBP 62 | 2H27, 6IDO, 1KU7, 1RIO, 5B7J, 5VXN, 5W43, 1LQ1, 3N97, 2Z33, 1ZLK, 2P7C, 1ZQ3, 1JE8, 1ZG5, 5CLV |

**Table S3. PDB IDs for native DBPs similar to designs.** DBP complexes with a TM-score greater than 0.65 which were computationally redesigned to see if a more naive native redesign strategy would recapitulate the *de novo* designed DBPs.

|  | **DBP048 + dsDNA* (PDB ID 8TAC)** |
| --- | --- |
| **Wavelength** | **0.9774** |
| **Resolution range** | **52 - 2.34 (2.424 - 2.34)** |
| **Space group** | **P 1 21 1** |
| **Unit cell** | **28.352 103.996 32.825 90 99.469 90** |
| **Total reflections** | **48632 (4751)** |
| **Unique reflections** | **7888 (772)** |
| **Multiplicity** | **6.2 (6.2)** |
| **Completeness (%)** | **99.57 (99.87)** |
| **Mean I/sigma(I)** | **5.38 (1.05)** |
| **Wilson B-factor** | **27.43** |
| **R-merge** | **0.1832 (0.2997)** |
| **R-meas** | **0.1998 (0.3272)** |
| **R-pim** | **0.07882 (0.1298)** |
| **CC1/2** | **0.979 (0.895)** |
| **CC*** | **0.995 (0.972)** |
| **Reflections used in refinement** | **7879 (771)** |
| **Reflections used for R-free** | **789 (77)** |
| **R-work** | **0.2441 (0.3196)** |
| **R-free** | **0.2793 (0.3653)** |
| **CC(work)** | **0.914 (0.529)** |
| **CC(free)** | **0.850 (0.511)** |
| **Number of non-hydrogen atoms** | **1442** |
| **macromolecules** | **1374** |
| **solvent** | **68** |
| **Protein residues** | **133** |
| **RMS(bonds)** | **0.003** |
| **RMS(angles)** | **0.58** |
| **Ramachandran favored (%)** | **96.90** |
| **Ramachandran allowed (%)** | **3.10** |
| **Ramachandran outliers (%)** | **0.00** |
| **Rotamer outliers (%)** | **5.81** |
| **Clashscore** | **12.66** |
| **Average B-factor** | **31.59** |
| **macromolecules** | **31.78** |
| **solvent** | **27.77** |
| **Number of TLS groups** | **9** |

**Table S4. Data collection and refinement statistics.** Statistics for the highest-resolution shell are shown in parentheses.
